# Self-Reactive B Cells in Artery Tertiary Lymphoid Organs Encode Pathogenic High Affinity Autoantibody in Atherosclerosis

**DOI:** 10.1101/2025.07.10.664251

**Authors:** Chuankai Zhang, Xi Zhang, Yi Ran, Zhihua Wang, Liping Li, Shu Wang, Junjie Zheng, Yixin Zhang, Ting Sun, Yutao Li, Shu Lu, Mingyang Hong, Zhe Ma, Sabine Steffens, Michael Hristov, Xavier Blanchet, Klaus Dornmair, Desheng Hu, Shibojyoti Lahiri, Axel Imhof, Nadja Sachs, Lars Maegdefessel, Jingyi Xiao, Jianning Zhang, Yan Wang, Hua Hong, Livia Habenicht, Christian Weber, Donato Santovito, Rachael Bashford-Rogers, Sarajo K. Mohanta, Klaus Ley, Andreas J.R. Habenicht, Changjun Yin

**Author notes:** These authors contributed equally (Chuankai Zhang, Xi Zhang, Yi Ran, Zhihua Wang). Senior author (Andreas J.R. Habenicht, Changjun Yin).

## Abstract

Artery tertiary lymphoid organs (ATLOs) emerge in atherosclerosis which is a chronic inflammatory artery disease with an autoimmune component. However, whether disease-relevant autoimmune B cells emerge in ATLOs and their impacts remains unknown. To map atherosclerosis-specific humoral autoimmunity and define its roles, we isolated germinal-center (GC) B cells from ATLOs and lymph-nodes, expression-cloned 60 autoantibodies and screened them for arterial wall reactivity. ATLO-derived autoantibodies markedly skewed to atherosclerosis-relevant autoantigens versus those of lymph-nodes. One ATLO GC-derived autoantibody bound to histone 2B (H2B) with high-affinity (∼25 nM). Moreover, vaccination with H2B or adoptive transfer of its cognate autoantibody markedly accelerated atherosclerosis suggesting that ATLOs fail to delete pathogenic high-affinity self-reactive B cells. In a human cohort, total circulating anti H2B antibody titers positively correlated with aortic and coronary artery calcification. We conclude that ATLOs harbor a dysregulated immune tolerance environment permissive for autoreactive B cells that express pathogenic autoantibodies driving atherosclerosis.

## INTRODUCTION

Atherosclerosis constitutes the major underlying disease pathology of heart attacks and strokes(Gisterå and Hansson, 2017; Libby, 2021) and is well recognized as a chronic inflammatory disease. However, its autoimmune disease etiology remains incompletely defined(Khan et al., 2024; Raposo-Gutiérrez et al., 2023; Sage et al., 2019; Wick et al., 2014). Previous data have shown atherosclerosis-associated autoreactive T cells(Khan et al., 2024; Wang et al., 2023) and autoreactive B cells(Raposo-Gutiérrez et al., 2023; Sage et al., 2019). Multiple autoantigens and autoantibodies have been identified: Monoclonal autoantibodies against oxidized low-density lipoprotein or aldehyde dehydrogenase 4A1(ALDH4A1) have atheroprotective effects and are of low or undefined affinity for their cognate antigen(Lorenzo et al., 2021; Nicolo et al., 2003; Que et al., 2018). oxLDL antibody titers negatively correlate with atherosclerotic cardiovascular disease (ASCVD) severity in humans(Shaw et al., 2000). Such autoantibodies may arise from peripheral naïve B cells that escaped central tolerance and express autoreactive B cell receptors (BCR), but do not trigger autoimmune disease(Shome et al., 2022). Polyclonal antibodies to β2GPI(Wang et al., 2019), GRP78(Crane et al., 2018) or CXCR3(Muller et al., 2023) promote atherosclerosis, but their affinities to the respective self-antigens have not been determined. Monoclonal antibodies to Hsp60(Foteinos et al., 2005) and Hsp70(Leng et al., 2010) were induced by vaccination and then shown to exacerbate atherosclerosis after adoptive transfer into recipient animals. None of these studies identified an endogenous autoantibody-autoantigen pair with defined specificity and affinity. Autoimmune diseases are defined by the Witebsky-Rose criteria(Rose and Bona, 1993), which require the identification of pathogenic autoantigens and their related high-affinity pathogenic autoantibodies that form autoimmune disease-causing pairs(Pisetsky, 2023; Rose and Bona, 1993). These criteria have not been met for atherosclerosis, and autoimmune disease societies do not list atherosclerosis as an autoimmune disease or autoimmune disease-related condition. Recent epidemiological studies have indicated that 19 autoimmune or autoimmune-related diseases confer risks for cardiovascular diseases including atherosclerosis(Conrad et al., 2022). Although the underlying mechanisms by which autoimmune disease confer risk for ASCVD remain unclear, breakdown of tolerance to self is a common denominator in many autoimmune diseases. Another prominent accelerator of ASCVD is immune checkpoint blockade in cancer treatment, which unleashes autoimmunity that manifests itself in exacerbated ASCVD(Poels et al., 2020; Thuny et al., 2022).

Tertiary lymphoid organs (TLOs) in general and artery tertiary lymphoid organs (ATLOs) in particular, unlike the stereotyped secondary lymphoid organs (SLOs), emerge exclusively in various diseased adult tissues in response to chronic inflammation such as in cancers, autoimmune diseases, transplant rejection, infectious diseases, and cardiovascular diseases(Bombardieri et al., 2017; Grabner et al., 2009; Hu et al., 2015b; Mohanta et al., 2022; Sato et al., 2023; Schumacher and Thommen, 2022; Yin et al., 2019). Curiously, clinical correlation studies of the incidence of TLOs have shown beneficial outcomes in most cancers and many infectious diseases(Schumacher and Thommen, 2022), but unfavorable outcomes in most autoimmune diseases, transplant rejection, and atherosclerosis(Corsiero et al., 2019; Hu et al., 2015b; Wieland et al., 2021). The functional impacts of TLOs on any of these diseases largely remain unknown. Earlier work has shown clonal expansion of T cells in ATLOs of mice with atherosclerosis indicating a T cell autoimmune component(Wang et al., 2023). ATLOs emerge in advanced murine and human atherosclerosis in disease-relevant segments of the arterial tree including coronary and carotid arteries and the abdominal aorta(Akhavanpoor et al., 2018; Grabner et al., 2009; Mohanta et al., 2022). Their advanced stages contain activated B cell follicles with germinal centers (GCs) and prominent plasma cell niches(Grabner et al., 2009; Srikakulapu et al., 2016). Thus, we suspected that ATLOs may participate in B cell autoreactivity and autoantibody affinity maturation(Srikakulapu et al., 2016). Autoreactive B cells are normally kept under control by highly efficient peripheral tolerance checkpoints(Brink and Phan, 2018; Vinuesa et al., 2009). When one or more tolerance checkpoints are compromised, autoimmune disease can result(Rose, 2017). Here, we focused on the humoral aspect of atherosclerosis by studying B cell and plasma cell transcriptomes and possible checkpoint defects. In GC B cells from ATLOs (and lymph-node controls), we determined immunoglobulin light and heavy chain pairs by combining single cell RNA sequencing (scRNA-Seq) and scBCR-seq in single cells. To test the relevance of these BCRs, we expression-cloned their corresponding antibodies, screened them for arterial wall and plaque reactivity and identified one pathogenic high-affinity monoclonal autoantibody and its autoantigen. Identifying relevant high-affinity autoantibodies that exacerbate atherosclerosis and their corresponding autoantigen(s) would strongly support an autoimmune etiology of atherosclerosis.

## RESULTS

### Dysregulated B and plasma cells in ATLOs

CD45^+^ leukocytes were FACS-sorted from pooled renal lymph nodes (RLN) of WT and Apoe^-/-^ mice and ATLOs from aged Apoe^-/-^ mice. After quality controls, 12,854 B cell transcriptomes - with matched scRNA-seq and scBCR-seq data - were retrieved. Uniform manifold approximation and projection (UMAP) algorithms identified 6 subsets (Fig.S1A,B,C): naїve B cells (antigen-inexperienced); newly activated B cells (Nac B cells); early GC B cells (non-dividing antigen-experienced GC B cells); late GC B cells (antigen-experienced GC B cells undergoing proliferation, somatic hypermutation (SHM), and affinity-driven selection); memory B cells (memB cells); and plasma cells (PCs, actively secreting antibody) (Fig.S1C,D). ATLOs harbored all prototypical B cell subsets of renal-lymph nodes (RLNs), but showed significantly higher percentages of early GC B cells, memB cells, and PC subsets vs both WT RLNs or Apoe^-/-^ RLNs (Fig.1A, Fig.S1B).

**Figure 1:**
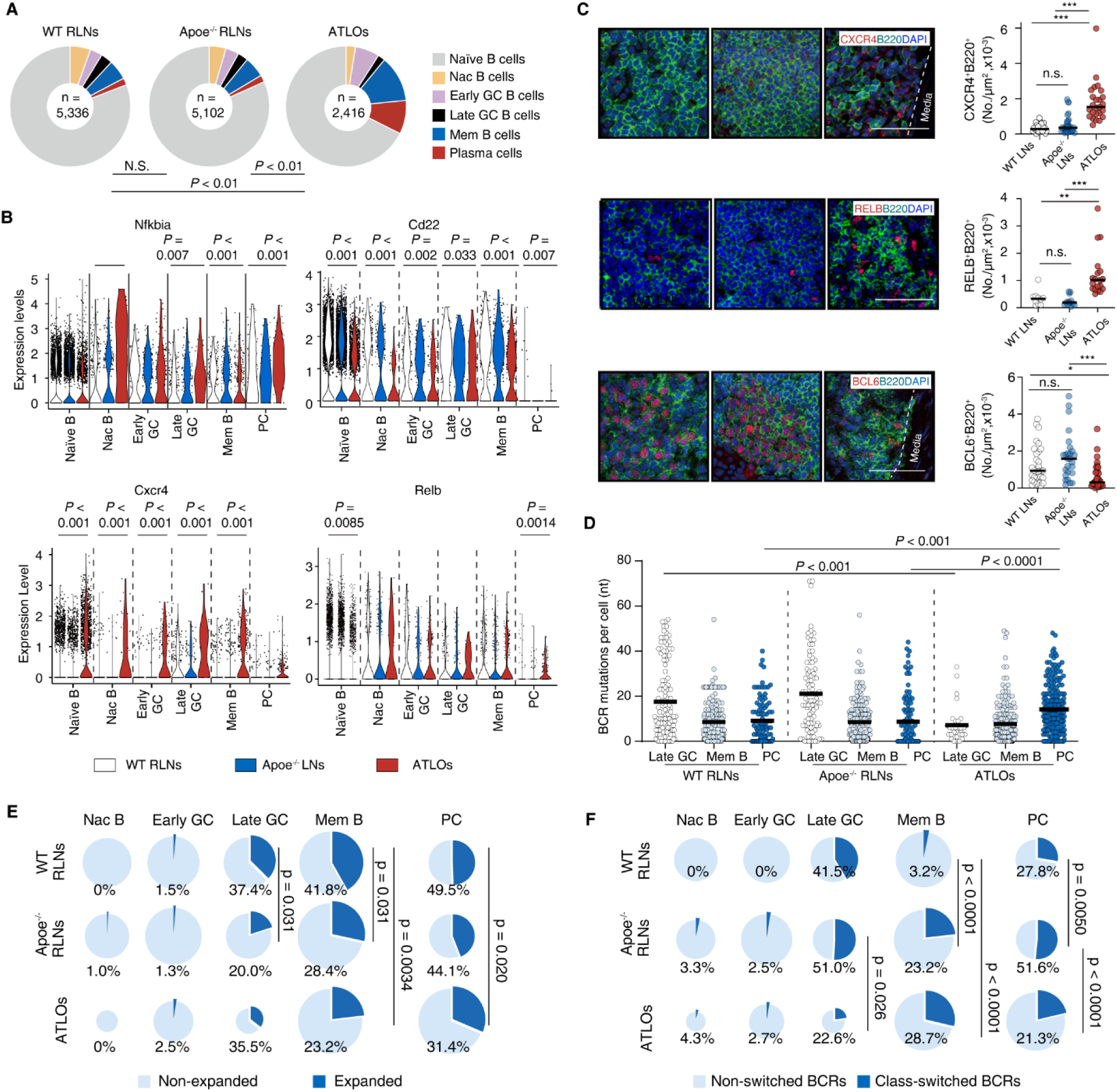
Delineation of scRNA-seq/scBCR-seq-paired B cell subsets reveals multilayered B cell tolerance dysfunction in ATLOs. (A) Relative composition of six B cell subtypes was determined based on their transcript profiles as shown in Figure S1; scRNA-seq identified six B cell subsets in each tissue according to their transcript profiles: naїve B cells, newly activated B cells (Nac B cells), early GC B cells, late GC B cells, memB cells and PCs; percentages of each B cell subtype in each tissue is indicated by different colors in the circle plots; the total number of B cells analyzed per tissue is shown in the inner circle. Statistical analysis was performed using the Chi-square test with Benjamini-Hochberg correction. (B) Selected transcripts in individual B cell subsets; Violin plots show B cell activation (Nfkbia, Relb), migration (CXCR4), the B cell inhibitory checkpoint (CD22) gene expression profiles in B cell subsets in different tissues. Each dot represents one single B cell. Statistical analysis was performed using the Kruskal-Wallis H Test with Benjamini-Hochberg correction. (C) Quantitative immunofluorescent analyses of key regulatory molecule-expressing cells in WT and Apoe^-/-^ RLNs and ATLOs; immunofluorescence followed by morphometry of CXCR4^+^ B cells and RelB^+^ B cells but lower numbers of BCL6^+^ B cells in B cell areas. Tissue sections were stained using anti-CXCR4, anti-RelB, anti-BCL6 and anti-B220 antibodies. Quantification of the number of positive cells/area per section of CXCR4^+^ B cells (migratory B cells), RelB^+^ B cells (activated B cells), and BCL6^+^ B cells (follicular B cells) within B220^+^ B cell areas were determined as described in Methods. Bars 100 μm. Each dot represents one field; 5-6 fields per mouse per tissue, n = 5 WT RLNs, 5 Apoe^-/-^ RLNs, and 5 ATLOs from 5 Apoe^-/-^ mice. (D) Number of somatic hypermutations (SHM) per BCR heavy chain; each dot represents one B cell. P values of two-group comparisons were determined by the Wilcoxon Rank Sum Test. (E) Pie charts show distinct ATLO traits of clonally expanded vs non-expanded B cell subtypes; sizes of circles represent the relative cell numbers within each tissue; blue indicates the percentage of clonally expanded B cells within each subset. (F) Percentages of class-switched BCRs in different B cell subsets WT and Apoe^-/-^ RLNs vs ATLOs; sizes of circles represent the relative cell numbers within each tissue; blue indicates the percentages of class switched B cell subtypes; G-H, Chi-square test with Benjamini-Hochberg correction was used to perform statistical analyses;

Differentially expressed gene (DEG) profiles in B cells showed major differences between ATLOs vs Apoe^-/-^ RLNs, but they were comparable between WT RLNs vs Apoe^-/-^ RLNs (Fig.S1E). All ATLO B cell subsets including naïve B cells, Nac B cells, early GC B cells, late GC B cells, and memB cells expressed higher levels of BCR activation-, T helper T cell-, and survival-related transcripts (Nfkbia, Relb, Cxcr4, Cd86), and lower levels of inhibitory- and apoptosis-related transcripts (Bcl6, Cd22, Fas) vs their counterparts in either WT RLNs or Apoe^-/-^ RLNs (Fig.S1F, Fig.1B). The high levels of CXCR4 and Relb proteins, and lower levels of BCL6 protein expressed by ATLO B cells were confirmed and quantified at the protein level (Fig.1C). ATLO late GC B cells, memB cells, and PCs showed distinct features of BCR SHM, including higher mutation numbers in ATLO PCs and lower mutation numbers in ATLO late GC B cells (Fig.1D). ATLO B cell subsets showed comparable BCR clonal expansion when compared to their counterparts in Apoe^-/-^ RLNs (Fig.1E). Since the highly mutated BCRs in PCs are mainly derived from GC responses(Victora and Nussenzweig, 2022), the significant higher number of BCR mutations in ATLO PCs did not support dysfunction of SHM mechanisms in ATLO GCs, but rather indicate that the GC-dependent PC differentiation pathway is enhanced in ATLOs. Although late GCs were enriched by class-switched GC B cells, the class-switching recombination (CSR) event occurs between T-B cell-cell interaction and SHM events in newly activated B cells and early GC B cells(Roco et al., 2019). In line with this model, the class-switched B cells were already detected in NacB cells and early GC B cells in Apoe^-/-^ RLNs and ATLOs (Fig.1F). Of note, the class-switched BCRs in late GC B cells and PCs are significantly lower than Apoe^-/-^ RLNs, and it remains unchanged in early GC B cells (Fig.1F), indicating CSR dysfunction occurs between early GC to late GC reactions in ATLOs. Moreover, ATLO PCs expressed higher Ig mRNA transcripts per cell vs PCs in WT and Apoe^-/-^ SLOs (Fig.S1G). Consistent with the markedly higher rate of Ig mRNA expression, ATLO PCs expressed significantly higher levels of unfolded protein response-related transcripts (Eif2ak3, Ppp1r5a) vs their counterparts in RLNs of both genotypes (Fig.S1H). ATLO PCs expressed higher levels of transcripts related to Ig secretion (Ighm), activation (Nfkbiz) and survival (Tnfrsf17) (Fig.S1I). This data indicated aberrant control of B cell/PC homeostasis in ATLOs (Fig.S1J). We next modeled T-B and myeloid-cell-B cell interaction networks within ATLOs vs those of WT and Apoe^-/-^ RLNs (Fig.S2A,B). ATLO B cells showed enhanced interactions with Lyve1^+^ resident-like macrophages, Trem2^high^ macrophages, CD8^+^ effector memory T cells, and γδ T cells vs their counterparts in RLNs (Fig.S2A,B). ATLO B cell transcriptomes were enriched in B cell survival, migration, and tolerance-regulating genes including TNFRSF13B (encoding TACI, a B cell survival factor), CCR5 and integrins (both controlling B cell trafficking), CD86 (associated with breakdown of self-tolerance) and CD80 (promoting B cell activation) (Fig.S2C,D,E). This data indicated that enhanced T-B and myeloid-B interactions during B cell trafficking through the pre-GC, GC, and post-GC compartments of ATLOs may contribute to B cell immune tolerance dysfunction in ATLOs (Fig.S2F).

### Dysregulated GC B cells in ATLOs

GCs arise in activated B cell follicles of lymphoid organs to organize B cell clonal expansion, SHM and BCR affinity maturation. Powerful tolerance mechanisms in GCs remove autoreactive B cells via multiple mechanisms including negative selection to eliminate emergence of autoreactive high-affinity pathogenic antibodies(Brink and Phan, 2018; Victora and Nussenzweig, 2022; Vinuesa et al., 2009). GCs have been reported to contain follicular dendritic cells (FDCs) in activated B cell follicles of TLOs in multiple disease conditions(Sato et al., 2023; Schumacher and Thommen, 2022; Victora and Nussenzweig, 2022). We observed considerable heterogeneity within ATLOs as defined by scattered T-B clusters (stage I) to advanced stages with CD35^+^ FDCs ranging from small patches of FDCs to larger GCs with well separated light zones (LZs, BCL6^+^CD35^+^) and dark zones (DZs, BCL6^+^CD35^-^) (stage 3) (Fig.2A-B). ATLO GCs were exclusively observed in Apoe^-/-^ mice with heavy atherosclerosis burdens(Grabner et al., 2009) (Fig.2B). This data indicates that ATLOs contain a range of immature (without GCs) to mature (with GCs) stages, reminiscent of those found in cancers(Zhang et al., 2024).

**Figure 2:**
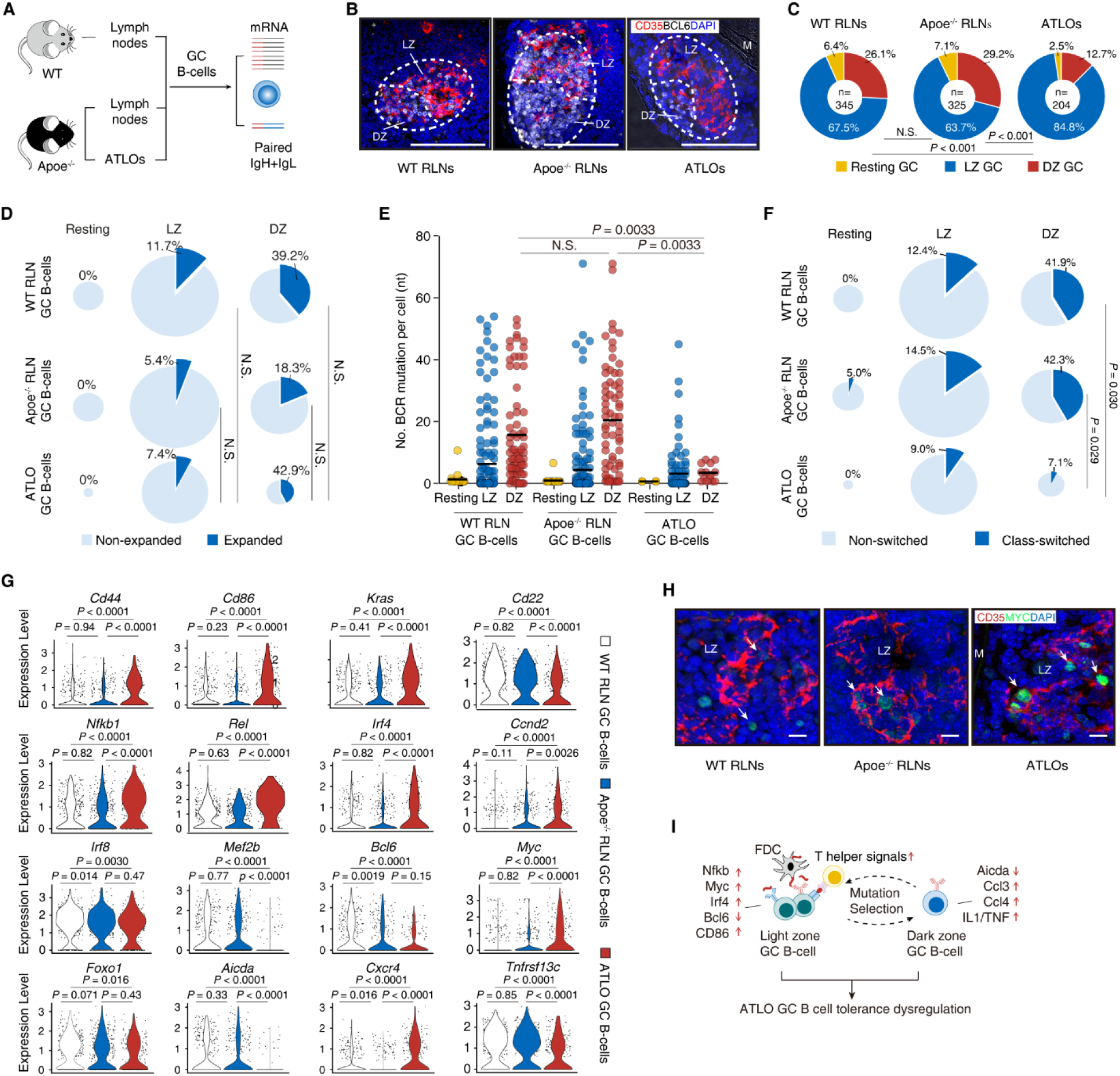
ATLO GCs reveal altered DZ-LZ structures and their B cell transcript signatures are dysregulated. (A) GC responses using pairing of scRNA-seq with scBCR-seq; pairing of scRNA-seq/scBCR-seq was performed as described in Methods. (B) Stage III ATLOs in activated B cell follicles accommodate separate LZs and DZs in GCs; adventitia segments adjacent of heavy atherosclerotic plaques were selected to stain FDC networks with anti-CD35 antibodies for LZs and B cell areas outside CD35^+^ networks were stained with anti-BCL6 antibodies for DZs. Data are representative of at least three independent experiments. (C) ATLOs reveal aberrant GC B cell composition; percentages of resting GC B cells, LZ GC B cells, and DZ GC B cells in WT RLNs, Apoe^-/-^ RLNs, and ATLOs as described in Methods. The total number of GC B cells analyzed in each tissue is shown in the inner circle. Chi-square test was used to compare the GC B cell subset compositions between two tissues. (D) ATLOs maintain expanded vs non-expanded B cell compositions comparable to WT GCs; percentages of clonally expanded vs non-expanded B cells in different GC B cell subsets in WT and Apoe^-/-^ RLNs and ATLOs. Chi-square post hoc test with Benjamini-Hochberg correction. (E) Quantification of SHM per antibody sequence imaged as Violin plots in ATLOs show dysfunction of BCR mutations per GC B cell; each dot represents one B cell. The P values of two-group comparisons was determined by Wilcoxon Rank Sum Test; P values of multiple-group comparisons was determined by Kruskal-Wallis H Test with Benjamini–Hochberg correction. (F) ATLOs reveal distorted CSR rates; percentages of CSR rates in different GC B cell subsets in different tissues. Chi-square test with Benjamini–Hochberg correction was used to perform statistical analysis. (G) Function-associated GC B cell gene expression profiles reveal exaggerated patterns; Violin plots show representative gene expression levels associated with GC B cell features: B cell activation (Cd44, Cd86), inhibitor of B cell activation (Cd22), BCR activation (Kras, Ccnd2), activation of costimulation (Nfkb1, Rel), master transcription regulators (Irf8, Mef2b, Myc, Bcl6), SHM signals (Foxo1, Aicda), B cell migration (Cxcr4) and survival signals (Tnfrsf13c/BAFFR). Each dot represents one B cell. Statistical analysis was performed using the Kruskal–Wallis H Test with Benjamini-Hochberg correction. (H) Higher numbers of myc^+^ GC B cellsin ATLOs; anti-CD35 antibodies and anti-Myc antibodies were used to stain WT RLN, Apoe^-/-^ RLN, and ATLO sections of aged WT and Apoe^-/-^ mice. (I) Schematic of aberrant features indicate ATLO GC B cell tolerance dysfunction.

We next explored ATLO GC B cells vs WT and Apoe^-/-^ RLN GCs at single cell resolution. We followed the algorithm to define the GC-derived B cell populations as reported previously (GC B cells obtained from WT mice vaccinated with 4-hydroxy-3-nitrophenylacetyl (NP) or ovalbumin, GSE154634)(Chen et al., 2021). GC B cells obtained from WT RLNs, Apoe^-/-^ RLNs and ATLOs were also separated into dark zone (DZ) GC B cells (Cxcr4^+^, Mki67^+^, Aicda^+^, Cdc20^+^; proliferating cells in the S-G2-M transition of the cell cycle), light zone (LZ) GC B cells (CD83^+^, Cxcr4^low^, CD74^+^), and resting GC B cells (Ccr6^+^, IgD^+^, Ly6d^+^) (Fig.S3A-E). WT and Apoe^-/-^ RLN GC B cells showed similar ratios of LZ to DZ GC B cells (Fig.2C), while ATLO GCs were distinct in their cellular composition with reduced DZs and increased LZs (Fig.2C).

To examine whether and ATLO GC B cells may be associated with functional changes, we examined clonal expansion, SHM, and class-switched BCRs of each GC subset. DZ GC B cells showed the highest levels of clonal expansion among all three GC B cell subsets in all three tissues (Fig.2D) consistent with the expression of high levels of proliferation-related genes as expected. There were no apparent differences between BCR clonal expansion among the three GC B cell subsets in WT RLNs, Apoe^-/-^ RLNs, and ATLOs (Fig.2D) indicating broad functional integrity of the GCs. In WT and Apoe^-/-^ RLNs, DZ GC B cells showed higher levels of SHM and isotype-switched BCRs when compared to LZ GC and resting GC B cells and there was no apparent difference between these two genotypes (Fig.2E,F). Interestingly, DZ GC B cells (but not LZ GC B cells) in ATLOs showed the lowest levels of BCR mutations and isotype-switched BCRs than their counterparts in WT RLNs and Apoe^-/-^ RLNs (Fig.2E,F). These dysregulated B cell responses indicate that ATLO GCs feature a dysregulated immune tolerance environment.

To understand the underlying mechanisms, we examined the gene expression by GC B cells obtained from different tissues. By two-group comparisons, the transcripts of GC B cells showed major overlaps between Apoe^-/-^ GC B cells vs WT GC B cells (Fig.S3F). However, ATLO GC B cells selectively upregulated B cell activation-(Fc receptors, NF-κB-related), interferon-related transcripts and downregulated B cell tolerance inhibitory (CD22) and SHM-related transcripts (Aicda) (Fig.S3F). Within GCs, B cells receive signals from antigen-engaged BCRs and costimulatory signals from FDCs and T follicular helper cells (TFH) cells(Brink and Phan, 2018; Victora and Nussenzweig, 2022; Vinuesa et al., 2009). We noticed significant changes of transcripts involved in multiple B cell-regulating pathways, including B cell activation (Cd44, Cd86), inhibitor of B cell activation (Cd22), BCR activation (Kras, Ccnd2), activation of costimulatory (Nfkb1, Rel), master transcription regulators (Irf4, Myc, Bcl6), positive regulation of Bcl6 (Irf8, Mef2b), BCR mutation and class switching (Foxo1, Aicda), B cell migration (Cxcr4), and B cell survival (Tnfrsf13c/BAFFR) (Fig.2G). In line with the changes at the transcript level, we observed apparent strong expression of Myc protein levels in ATLO LZ GC B-cell regions (Fig.2H). This data revealed that ATLO GC B cells receive exaggerated signals from T helper cells which may indicate tolerance dysfunction*(34)*.

Next, we performed gene set enrichment analyses (GSEA) of ATLO LZ GC B cells and DZ GC B cells vs their counterparts in WT RLN GCs and Apoe^-/-^ RLN GCs (Fig.S4A-D). ATLO LZ GC B cells were positively enriched in GO terms of CD4 T cell activation, DC cell differentiation, and tolerance induction and inhibition, and negatively enriched in GO terms in DNA replication and the BCR receptor signaling pathway (Fig.S4B,C). Interestingly, ATLO DZ GC B cells were positively enriched in GO terms of TNF and IL-1β production, and negatively enriched in the somatic hypermutation-related GO term (Fig.S4B,D). Moreover, ATLO LZ GC B cells expressed high levels of T helper cell differentiation (Nfkbid, Irf4, Stat3)-, tolerance breakdown (Cd86, Ccr7)-, and tolerance enhancement (Cd274, Tnfaip3, Tgfb1, Cblb)-related transcripts (Fig.S4E,F). ATLO DZ-GC B cells expressed high levels of members of the TNF family and IL-1β production- and chemokine-mediated signaling pathway-related transcripts (i.e., Il1b, Ccl3, Ccl4, Ccl6, Ccl9) (Fig.S4E,F). This data suggests that the different B cell signals in LZ and DZ GC B cells (Fig.2I) may determine distinct B cell fates vs their counterparts in RLNs of both genotypes and that these phenotypes may contribute to the aberrant GC structure of ATLOs.

### ATLO GC-derived autoantibodies skew to atherosclerosis-relevant autoantigens

To study the functional impact of the apparent tolerance dysfunction of ATLO GC B cell responses, we cloned the paired IgH/IgL chains specifically from single GC B cells that had been isolated from WT RLNs, Apoe^-/-^ RLNs, and ATLOs, and generated the recombinant antibodies *in vitro*, followed by antibody functional assays (Fig.3A). We obtained 30 antibodies from RLNs (13 from WT RLN GCs, 17 from Apoe^-/-^ RLN GCs) and 30 from Apoe^-/-^ ATLO GCs. None of the 60 expression-cloned antibodies shared H or L CDR3 sequences (Table.S1). First, we determined whether ATLO GC B cells undergo affinity maturation, which is associated with reduced polyreactivity. To understand poly-vs oligoreactivity, we measured antibody binding to three structurally unrelated antigens, i.e. insulin, double-stranded DNA (dsDNA), and lipopolysaccharide (LPS), all representing self-antigens associated with low-affinity-binding antibodies(Notkins, 2004). 54% of WT RLN GC-derived antibodies and 41% of Apoe^-/-^ RLN GC-derived antibodies were polyreactive (binding at least 2 of the 3 antigens, Fig.3B, Tabl.S1). By contrast, only 10% of ATLO GC-derived antibodies were polyreactive (Fig.3B, Tabl.S1). Decreasing antibody polyreactivity in GC reactions is expected to be associated with increasing antigen specificity(Notkins, 2004).

**Figure 3:**
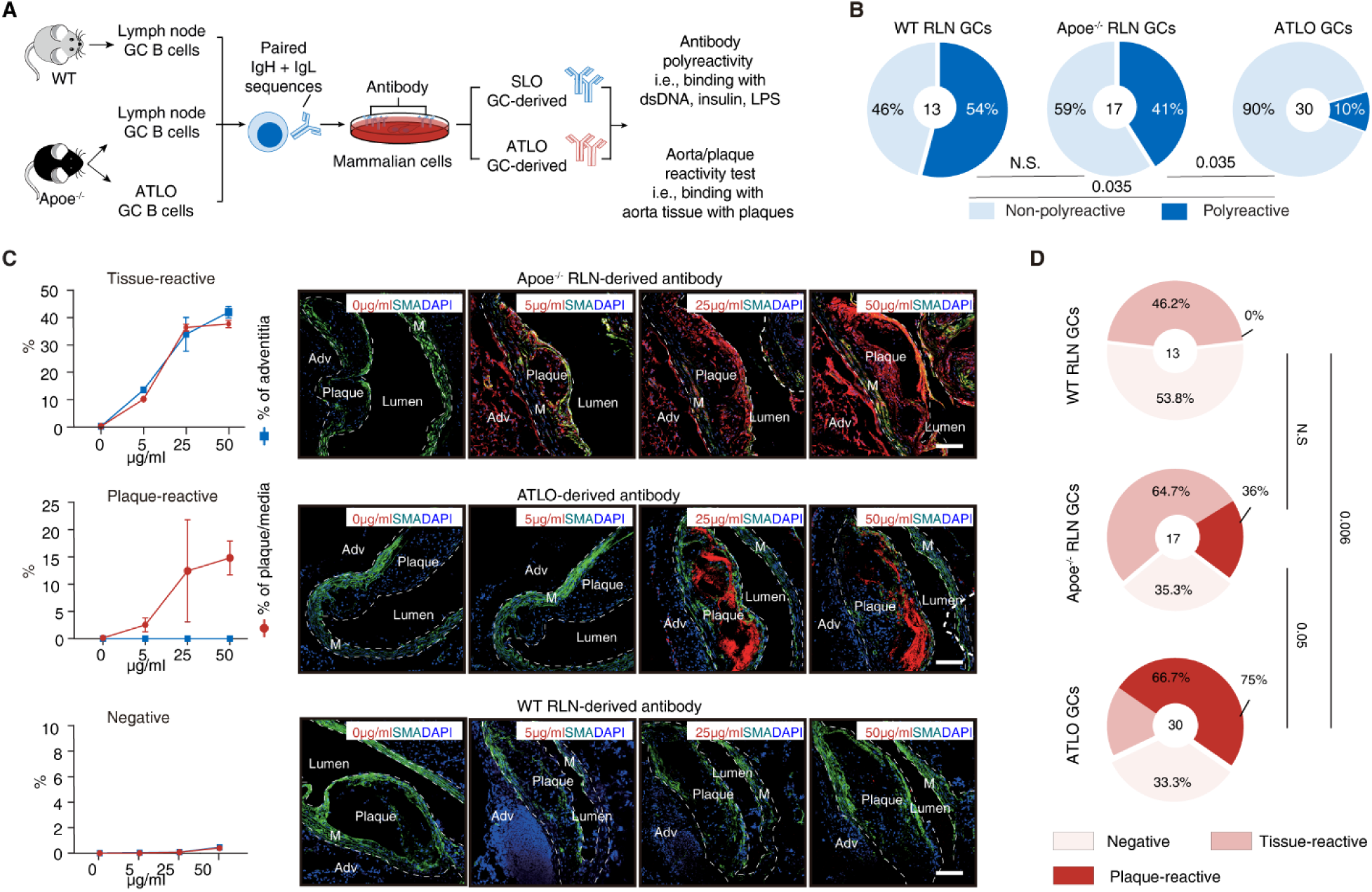
ATLO GCs generate diseased arterial wall-specific autoantibodies with high rates of plaque reactivity. (A) Schematic view of experimental design of sequencing, cloning, expression and characterization of GC B cell-derived autoantibodies. The variable regions of paired Ig heavy (IgH) + Ig light (IgL) chains were sequenced followed by cloning of the paired sequences into expression plasmids; following transfection into and expression by cultured cells, antibodies were purified from the culture supernatant. Antibodies were examined for immunoreactivities towards tissues and broad autoantigens, i.e. double-stranded DNA; insulin; lipopolysaccharide; antibodies were screened for reactivity towards aorta/plaque tissues by IF staining. (B) ATLOs selectively show markedly reduced polyreactivity; antibody polyreactivity against ≥ 2 of three structurally unrelated antigens: LPS, insulin, dsRNA were defined as polyreactive. Quantification of polyreactive antibodies in WT RLN GC-, Apoe^-/-^ RLN GC-, and ATLO GC-derived antibodies. The number of antibodies examined are listed in the inner circle. Two-sided Fisher’s exact test. (C) IF staining of three different patterns of antibody in murine atherosclerotic plaques. Different antibody concentrations (0, 5, 25, 50 μg/ml) were used to stain serial sections of arteries. Three antibody staining patterns are shown; i. no staining (aorta-negative) of all tested antibody concentrations; ii. stains positive for atherosclerotic plaques, media, adjacent adipose and connective tissues (aorta-reactive); iii. stains positive for plaques and media (plaque-reactive); bars 100 μm; (D) Quantification of different antibody patterns in RLNs vs ATLO GC-derived antibodies; ATLO GC-derived antibodies show high plaque-reactivity. The number of antibodies is listed in the inner circle. Chi-Square test;

Next, we tested all ATLO GC-derived autoantibodies and RLN GC-derived autoantibodies for binding to arterial walls including atherosclerotic lesions. For this purpose, we stained atherosclerotic mouse aorta tissues including plaque, media, adventitia and adjacent adipose tissue (Fig. 3C). By immunofluorescence staining, we observed that 46% (6/13) of WT RLN GC-derived antibodies, 65% (11/17) of Apoe^-/-^ RLN GC-derived antibodies, and 67% (20/30) of ATLO GC-derived antibodies recognized the diseased aorta and adjacent adipose tissue (referred to hereafter as “tissue-reactive antibodies”, Fig.3C,D, Tabl.S1). A subset of tissue-reactive antibodies (36%, 4/11 Apoe^-/-^ RLN GC B cells and 75%, 15/20 ATLO GC B cells) specifically recognized atherosclerotic plaques (“plaque-reactive”) (Fig.3C,D, Tabl.S1). In contrast, none of the tissue-reactive antibodies (0/6) obtained from WT RLN GCs were plaque-reactive (Fig.3D). The significantly higher levels of plaque-reactive antibodies in ATLO GCs (Fig.3D) suggests that ATLO GCs generated self-reactive B cells that were skewed towards plaque reactivity. This data indicated that ATLO GCs are functionally distinct when compared to their brethren in SLO GCs.

### ATLO GCs encode high-affinity autoantibody in atherosclerosis

To investigate the specificity of the ATLO GC-derived antibody responses, we focused on mAb A6 derived from ATLO GC B cells (Fig.4A), because A6 showed strong plaque-reactivity in arteries of aged Apoe^-/-^ mice (Tabl.S1) with a nuclear binding pattern (Fig.4B) and also showed reactivity to nuclei in human atherosclerotic plaques (Fig.4C). mAb A6 recognized a ∼10-15 KDa antigen in mouse atherosclerotic tissues (Fig.4D). Pull-down of the A6 candidate autoantigens from murine aorta extracts using immunoprecipitation followed by mass spectrometry (Fig.4E,F) identified histone-2b (H2B) as the cognate autoantigen for ATLO-GC-derived mAb A6 (Fig.4E,F). H2B is identical at the amino acid sequence level between mice and humans, explaining the mouse-human cross-immune reactivity of mAb A6. H2B is a known autoantigen involved in several clinically important diseases, including systemic lupus erythematosus(Kaul et al., 2016). However, its participation in atherosclerosis autoreactivity has not been reported. We validated the binding features of recombinant H2B to A6 by multiple methods, including western blot, surface plasmon interferometry (SPR) and biolayer interferometry (BLi) (Fig.4G,H).

**Figure 4:**
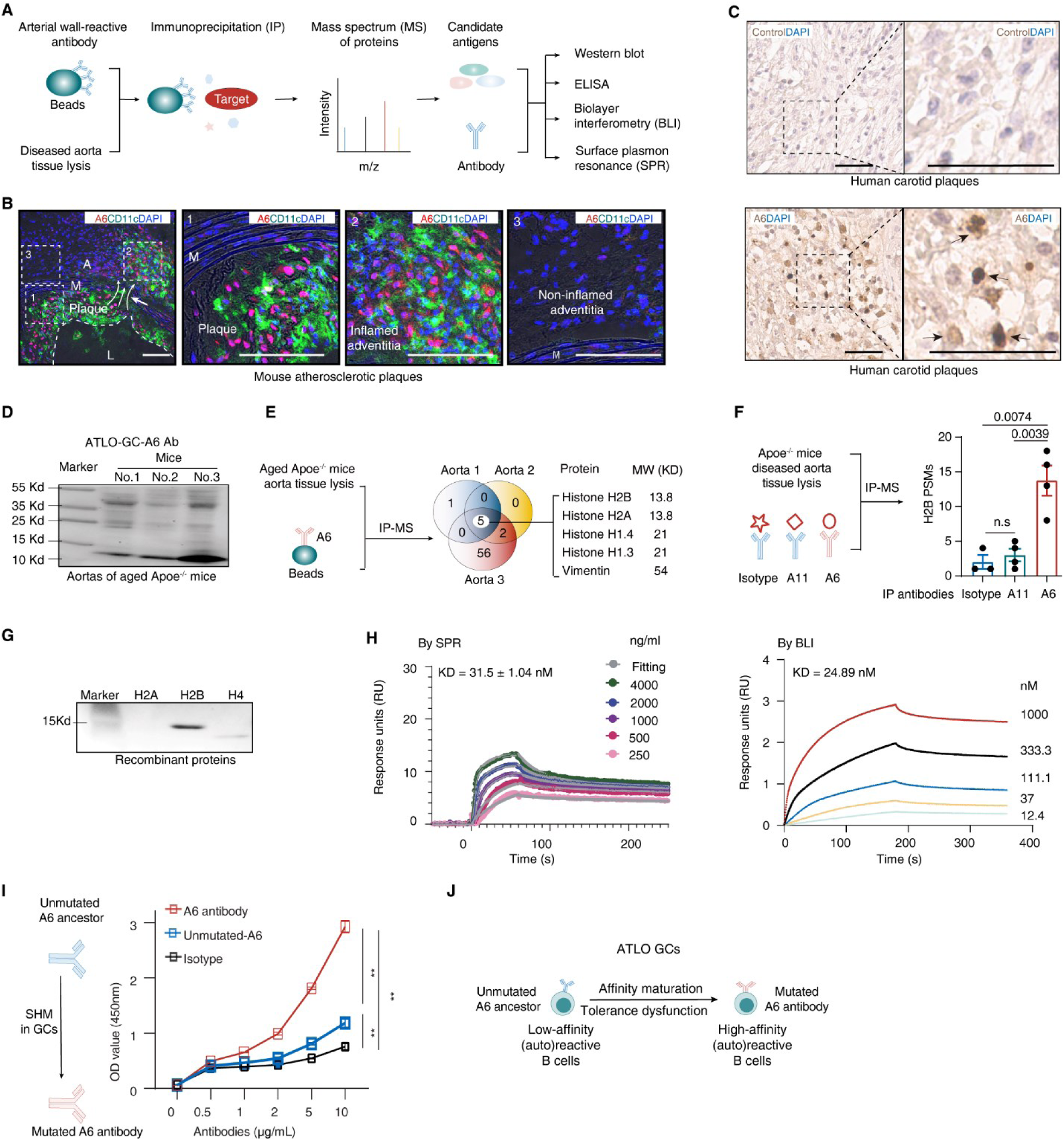
ATLO GCs give rise to a high-affinity autoantibody clone (termed A6) directed against histone 2b (H2B) (A) Identification of the cognate autoantigen of A6. Schematic view shows the workflow to identify atherosclerosis autoantigens using diseased artery-detecting antibodies. A6-loaded protein A beads were incubated with tissue lysates prepared from diseased mouse arteries followed by elution of proteins from the beads to undergo autoantigen screening of candidates by mass spectrometry (MS) analyses. Several autoantigen-antibody interaction assays were used to confirm protein-protein interaction in vitro including western blots, ELISA, biolayer interferometry (BLI) and surface plasmon resonance (SPR) analyses using various expression-cloned autoantibodies derived from RLNs and ATLO GCs. (B) A6 reacts with mouse aorta-associated atherosclerotic plaques; co-staining of A6, macrophages/DCs and nuclei in Apoe^-/-^ atherosclerosis plaque sections. A6 (red), CD11c (green) and nuclei (DAPI, blue). Bars 100 μm. (C) A6 reacts with human aorta or carotid artery atherosclerotic plaques; staining of A6 in human carotid atherosclerosis plaques by IHC. Bars 50 µm. (D) A6 identifies a 10-15 kDa protein in mouse diseased artery lysates; western blot (WB) shows A6 staining of aorta tissues obtained from 3 individual aged Apoe^-/-^ mice. (E) Immunoprecipitation (IP) combined with MS analyses to identify autoantigen candidates. The ATLO GC-derived A6 antibody was used as a bait to capture autoantigens from protein lysates of diseased aorta of 78-weeks aged Apoe^-/-^ mice. Beads without primary antibody was used as control. Label-free quantitation (LFQ) values of A6 were compared to controls. A6 antibody-enriched candidates are shown as a Venn diagram from three aorta samples from three individual aged Apoe^-/-^ mice. The molecular weight (MW) of each candidate protein is shown. (F) A6 antibody captured histone 2b (H2B) protein in the diseased aorta. The ATLO GC-derived A6 antibody was used as a bait to capture autoantigens from protein lysates of diseased aorta of 32-weeks Apoe^-/-^ mice. An isotype control antibody and the ATLO-GC-derived A11 antibody were used as controls. The peptide-spectrum matches (PSM) values of H2B are shown. Each dot represents one individual biological repeat. Three 32-weeks old Apoe^-/-^ aortas were combined as one sample. (G) A6 binding to recombinant H2B by WB; recombinant H2a, H2b, H4 were used for comparison. (H) A6 reacts with human recombinant H2B at high-affinity; binding affinity of A6 to human recombinant H2B by SPR and BLI analyses. (I) A6 acquires high-affinity binding to the H2B autoantigens via SHM from its unmutated ancestor; binding of ATLO GC-derived A6 antibody to H2B determined by ELISA: Mutated A6 (red line), the unmutated parent of A6 (unmutated-A6, blue line), and the isotype control antibody (black line). (J) Summary figure: A6 acquires high-affinity binding to H2B via ATLO GC responses indicating that negative selection of GC autoreactive BCRs to prevent high-affinity autoantibody generation is compromised in ATLOs.

Under homeostatic conditions, GC negative selection mechanisms of self-reactive BCRs are indispensable for the maintenance of self-tolerance to prevent generation of high-affinity pathogenic autoantibodies(Brink and Phan, 2018; Victora and Nussenzweig, 2022; Vinuesa et al., 2009). Here, we found that A6 bound recombinant H2B with high affinity (25∼30 nM, Fig.4H), suggesting that ATLO GCs allow the survival of B cells expressing high-affinity self (H2B)-reactive BCRs. To directly test the impact of the ATLO GC reaction on affinity maturation, we generated a recombinant germline unmutated A6 (unmutated-A6 mAb, Fig.4I). The unmutated-A6 had much lower binding affinity to H2B than mAb A6 (Fig.4I), demonstrating that A6 had acquired high-affinity binding for H2B during ATLO GC SHM and affinity maturation. That B cells with BCRs encoding high-affinity mAb A6 to the self-antigen H2B were found in ATLO GCs indicates dysfunction of negative selection during their GC reactions (Fig.4J).

### Humoral H2B autoimmunity drives atherosclerosis in mice

We next examined the functional impact of mAb A6 and the H2B autoantigen on atherosclerosis. We first determined the total anti H2B antibody titers in WT and Apoe^-/-^ mice during aging (Fig.5A). Serum anti H2B IgG1 antibody (Fig.5A) titers increased in Apoe^-/-^ mice during atherosclerosis progression vs age-matched WT controls, establishing an association between the H2B autoimmune humoral responses and atherosclerosis. To study the potential pathogenicity of the H2B autoimmune response in atherosclerosis, we established an H2B-induced vaccination protocol in Apoe^-/-^ mice. A mixture of recombinant H2B antigen and adjuvant (termed here as H2B vaccine) was given to 8 weeks old Apoe^-/-^ mice, i.e. before major atherosclerosis plaques emerge in this mouse model followed by a boost 4 weeks after the first injection. 6 months after the first injection, mice were sacrificed to determine atherosclerosis plaque loads (Fig.5B). Vaccinating with H2B significantly increased atherosclerotic plaque sizes in the whole aorta (Fig.5B) including aortic roots (Fig.S5A). Serum anti H2B-specific antibody responses were determined by ELISA. The H2B vaccine induced long-lasting anti H2B-specific antibody responses (Fig.5C). Significantly higher levels of T cell dependent IgG1-specific H2B antibody titers were observed in the H2B vaccine group when compared to controls (Fig.5D). Body weight and blood lipid levels remained unchanged (Fig.S5B,C). This data demonstrates that H2B autoimmune reactivity promotes atherosclerosis by inducing H2B humoral antibody responses independent from lipid levels.

**Figure 5:**
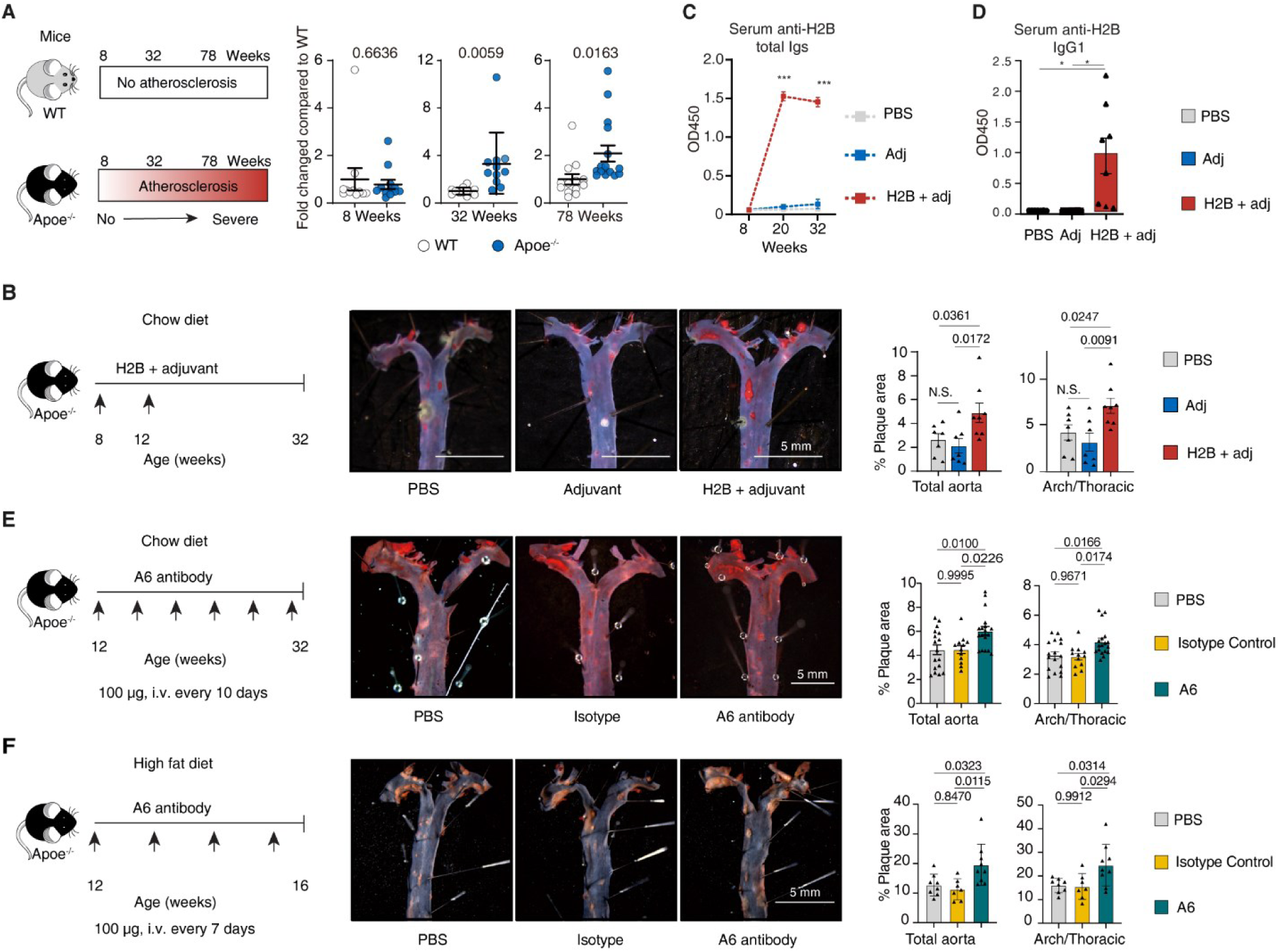
Atherosclerosis pathogenicity of the H2B-A6 pair. (A) Anti-H2B autoantibody titers increase in Apoe^-/-^ mice during aging. Anti-H2B IgG1 serum antibody titers of WT and Apoe^-/-^ mice during aging were examined by ELISA; each dot represents one mouse. n= 8-weeks WT (n = 11), Apoe^-/-^ (n = 12); 32-weeks WT (n = 12), Apoe^-/-^ (n = 11); 78-weeks WT (n = 13), Apoe^-/-^ (n = 16). Two-tailed student T test. (B) H2B vaccination promotes atherosclerosis. Vaccination using recombinant H2B in young Apoe^-/-^ mice. Apoe^-/-^ mice were vaccinated with recombinant H2B with adjuvant (termed as H2B vaccine), adjuvant alone, or PBS alone at the age of 8 weeks with a boost at the age of 12 weeks. The experiment was completed at 32 weeks. Blood was collected at different time points before and after the injection; *en face* staining for whole aorta. Atherosclerotic plaques were quantified, as described in Methods. Scale bar 5 mm. for PBS control (n = 7 mice), adjuvant control (n = 7), H2B + adjuvant (n = 8). (C-D) H2B vaccination induced long-lasting T cell dependent autoantibody responses in Apoe^-/-^ mice. Serum anti-H2B total antibody and anti-H2B IgG1 antibody titers were determined by ELISA. n = 7 PBS, 7 adjuvant, and 8 H2B + adjuvant. (E) A6 antibody transfer accelerated atherosclerosis in chow diet-fed Apoe^-/-^ mice. 100 µg A6 antibody was transferred via intraperitoneal (i.p.) injection at 10 weeks, and injections were repeated every 10 days. The experiment was completed at 32 weeks. Blood was collected at multiple time points before and after the injection; *en face* staining for whole aorta. Atherosclerotic plaques were quantified, as described in Methods. Scale bar 5 mm for PBS control (n = 17 mice), isotype antibody control (n = 12), A6 antibody (n = 18). (F) A6 antibody transfer accelerated atherosclerosis in high fat diet-fed Apoe^-/-^ mice. 100 µg A6 antibody was transferred via i.p. injection at 10 weeks, and injections were repeated every 7 days. The experiment was completed at 14 weeks. Blood was collected at several time points before and after the injection; *en face* staining for whole aorta. Atherosclerotic plaques were quantified, as described in Methods. Scale bar 5 mm for PBS control (n = 8 mice), irrelevant isotype antibody control (n = 7), A6 antibody (n = 9).

To directly test the pathogenicity of a high-affinity autoantibody (i.e., mAb A6), we transferred A6 antibody to two groups of young Apoe^-/-^ mice maintained on either chow or high-fat-diet, respectively (Fig.5E,F). A6 antibody transfer significantly increased atherosclerosis plaque loads under both dietary regimens compared to saline or irrelevant isotype controls (Fig.5E,F). Mouse body weights and blood lipid levels were not affected by A6 transfer (Fig.S5D,E). This data shows that high-affinity A6 autoantibody is pathogenic and promotes atherosclerosis in mice independent of lipids.

### Serum anti-H2B antibodies are associated with aortic calcification in humans

Finally, we tested whether the humoral autoimmune response to H2B is associated with atherosclerosis in humans (Fig. 6A). Serum anti H2B antibody titers were determined in a cross-sectional cohort of 495 individuals representative of the general population in China aged between 30 - 70 years who underwent chest computed tomography (CT) scans as part of a routine medical preventive screening program (Fig.6A, demographic and clinical characteristics in Table. S2). Higher serum anti-H2B antibody titers were associated with the prevalence of thoracic aortic wall calcification (TAC). TAC is commonly found in chest CT screening and is closely related to systemic atherosclerosis, morbidity and mortality(Desai et al., 2018). The association between TAC and anti H2B antibody titers persisted (P=0.023) after adjustments for age, male sex, LDL cholesterol, systolic blood pressure, HbA1c (reflecting long-term plasma glucose homeostasis), and estimated glomerular filtration rate (eGFR, reflects kidney function) by logistic regression (Fig.6B). This data is consistent with the possibility that humoral autoimmunity to H2B contributes to atherosclerosis in humans (Fig.6C).

**Figure 6:**
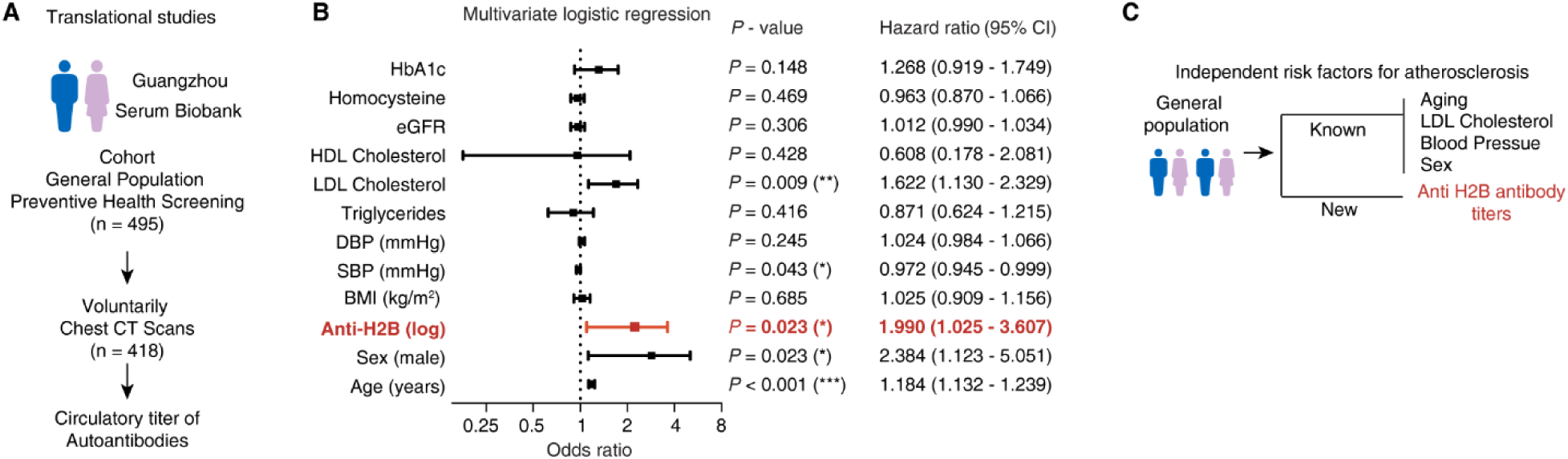
Serum anti-H2B antibody titers is associated with aorta calcification. (A) study design of the clinical cohort. A cross-sectional cohort of 495 individuals of the general population aged between 30 - 70 years who underwent routine medical screening of preventive health care was studied. Exclusion criteria included patients with previously diagnosed cardiovascular diseases, strokes, autoimmune diseases requiring systemic treatment, active malignancies, and severe kidney or liver dysfunctions. 84.4% (418 out of 495) cases voluntarily underwent chest 3-dimentional computed tomography (CT) scans as part of the preventive health screening program. Serum anti-H2B antibody titers were examined. (B) Serum anti-H2B antibody titers is an independent risk factor for aorta and coronary artery calcification. Anti-H2B antibody titers were logarithmically transformed before entering the model to approximate Gaussian distribution. Odds ratios and 95% confidence intervals (CI) are calculated by binary logistic regression with simultaneous entry of all variables (Cox & Snell *R*^2^ 0.296, *P* <0.001). eGFR, estimated glomerular filtration rate; HbA1c, glycated hemoglobin A1c; DBP, diastolic blood pressure; SBP, systolic blood pressure; BMI, body mass index; (C) Serum anti-H2B antibody is an independent risk factor for atherosclerosis.

## DISCUSSION

Here, we identify the high affinity mAb A6 and its cognate self-autoantigen H2B as an endogenous atherosclerosis-relevant autoantibody-autoantigen pair. Vaccinating Apoe^-/-^ mice with H2B or adoptive transfer of A6 both significantly exacerbated atherosclerosis, and antibody titers to H2B were positively correlated with thoracic aortic wall and coronary artery calcification, a hallmark of atherosclerosis in humans. mAb A6 was cloned from paired heavy and light chain BCR sequences from GC B cells in ATLOs, suggesting that ATLOs are lymphoid tissues with a broken tolerance to self (Fig.S6). This is further supported by significant transcriptomic differences between ATLO GC B cells and GC B cells in LNs, i.e., Bcl6, Cxcr4, and Nfkb. Unlike other B cell subtypes GC B cells do not circulate(Victora and Nussenzweig, 2022), so there is likely no exchange between lymph-nodes and ATLO GC B cell populations, further supporting the notion that the unique immune environment in ATLOs allows the expansion, somatic hypermutation, and affinity maturation of self-specific autoreactive B cells (Fig.S6).

Autoantibodies in atherosclerotic mice and humans have been described for decades(Raposo-Gutiérrez et al., 2023). Antibodies against oxLDL or ALDH4A1 have atheroprotective effects and are of low or undefined affinity for their cognate antigens(Lorenzo et al., 2021; Nicolo et al., 2003; Que et al., 2018). oxLDL antibody titers negatively correlate with atherosclerotic cardiovascular disease (ASCVD) severity in humans (Shaw et al., 2000), suggesting that they may be atheroprotective, not atherogenic. On the other hand, polyclonal antibodies to β2GPI(Wang et al., 2019), GRP78(Crane et al., 2018) or CXCR3(Muller et al., 2023) promote atherosclerosis, but their affinities to the respective self-antigens have not been determined. Adoptive transfer of monoclonal antibodies to Hsp60(Foteinos et al., 2005) or Hsp70(Leng et al., 2010) induced by vaccination was shown to exacerbate atherosclerosis in recipient animals. Notably, none of these previous studies identified an endogenous autoantibody-autoantigen pair with defined specificity and affinity, which sets our discovery apart. First, we used an unbiased approach to clone the candidate mAbs from BCR heavy and light chain sequences. Second, we screened for plaque reactivity and against general tissue reactivity. Third, because we know the origin of each B cell based on the experimental design and its transcriptome, we can pinpoint the origin of the A6 high affinity autoantibody to H2B to ATLO GCs (Fig.S6). ATLOs are atherosclerosis-specific in that they are not found in healthy arteries (Wang et al., 2023).

As little is known about the specific disease impact of any TLO in any disease(Sato et al., 2023; Schumacher and Thommen, 2022), mechanisms of ATLO tolerance dysfunction in atherosclerosis identified here may apply to other TLO-associated diseases(Fridman et al., 2022; Mohanta et al., 2014; Sato et al., 2023; Schumacher and Thommen, 2022; Zhang et al., 2024). It would be of considerable interest to compare mechanisms of tolerance control in TLO-associated conditions with opposite prognostic disease outcomes such as in most types of cancers with protective vs those with detrimental outcomes as in autoimmune diseases and atherosclerosis(Grabner et al., 2009; Sato et al., 2023; Sautes-Fridman et al., 2019; Zhang et al., 2024). We hypothesize that TLOs of diverse diseases containing GCs in activated B-cell follicles and separate T-cell areas may be similar to some ATLO tolerance traits described here: All form in response to chronic inflammation close to or within the diseased target tissue; all are home to a large innate inflammatory immune cell compartment; all have separate T cell and B cell areas and many are home to FDCs in addition to developing structures to promote T cell and B cell homing(Grabner et al., 2009; Sato et al., 2023; Schumacher and Thommen, 2022). Though we suspect similarities of tolerance dysfunction in all TLO-related diseases, there may also be significant dissimilarities: These may be related to the functionally T cell and/or B cell subtypes as well as the nature of the autoantigens differentially driving these diseases. For example, in atherosclerosis, CD8^+^ T cells may promote atherosclerosis while cancer autoantigen-specific CD8^+^ T cells may be beneficial to eliminate cancer cells. To distinguish between these possibilities it will be important to develop antigen- and autoantigen-specific next generation therapeutics.

Our discovery of a pathogenic autoantigen-autoantibody pair in ATLOs represents the first identification of such a pair in any TLO of any organ or disease. Thus, we establish a new paradigm: that the immune environment in TLOs is uniquely suited to initiate autoimmune response and different from the immune environment in LNs. First, TLOs have no capsule. Whereas LNs have significant anatomical and biological structures to facilitate entry of antigen presenting cells (APCs), it is likely that more and perhaps different APCs can access TLOs. The nature of the atherosclerosis-relevant APCs is not known, but we speculate that they may be some kind of dendritic cells. Second, TLOs are enriched with inflammatory macrophages and it is unknown how this trait may alter the immune environment in ATLOs or other TLOs. Since our approach is, in principle, applicable to other diseases and disease models, high-affinity autoantibodies can perhaps be cloned from cancer TLOs or any other TLO in any disease. Recently, TLO-derived antibodies directed against exogenous autoantigens have been shown to be involved in controlling certain cancers(Wieland et al., 2021; Zhang et al., 2024).

Breaking of tolerance to self has previously been shown in CD4^+^ and CD8^+^ T cells in a mouse model of atherosclerosis(Wang et al., 2023) and in human plaques(Depuydt et al., 2023). This begs the question whether loss of T cell tolerance precedes loss of GC B cell tolerance. An argument in favor of this interpretation is that CD4^+^ TFH cells are required for most isotype switching and affinity maturation of GC B cells. Once T cell tolerance is lost, self-reactive CD4^+^ T cells may differentiate into TFH cells and support affinity maturation. In a large number of T cells, tolerance to self is encoded in the transcriptome of the peripheral naïve CD4^+^ repertoire(Nettersheim et al., 2025). Naïve self-reactive CD4^+^ T cells express more CD73 and PD1 than naïve foreign-reactive CD4^+^ T cells, and blocking these molecules unleashes a CD4^+^ T cell-triggered autoimmune response. A second pathway to breaking tolerance may be the conversion from T_reg_ cells to exT_reg_ cells(Freuchet et al., 2023; Sharma et al., 2009). Interestingly, a recent study in human tonsil-derived organoids shows that knocking out FoxP3, the T_reg_-defining transcription factor, in CD4^+^ T cells allows the formation of high-affinity autoantibodies(Chen et al., 2025). It remains to be seen whether a loss of FoxP3 in CD4^+^ T cells in ATLOs(Wang et al., 2023) may be sufficient to allow autoreactive B cells to develop, proliferate and become antibody-secreting PCs.

H2B is not atherosclerosis-specific, but it is a known autoantigen in other classic autoimmune diseases(Kaul et al., 2016). Nuclear DNA is normally tightly coiled around histones. Our findings suggest that H2B may become “visible” to the immune system in atherosclerotic lesions and ATLOs. This could be related to neutrophil extracellular traps (NETs) that are known to occur in atherosclerosis(Silvestre-Roig et al., 2019). Since NETs consist of DNA, histones and other proteins, H2B may become exposed in NETs. However, multiple other means are known to lead to humoral autoimmune responses against nuclear proteins.

In conclusion, the discovery of pathogenic high affinity autoantibodies to H2B provides an important element supporting the autoimmune nature of clinically significant atherosclerosis. That these autoantibodies come from ATLOs suggest that ATLOs are functionally important to impact autoimmunity in atherosclerosis (Fig.S6).

### Limitations of the study

Limitations of this study include that it was performed in a mouse model of atherosclerosis, which does not match all aspects of human atherosclerosis. However, the discovery that H2B antibody titers are positively correlated with atherosclerosis in humans suggest that our findings are translatable. Another limitation is that our study was not designed or powered to discover sex differences. Autoimmune diseases are more common in women than men(Pisetsky, 2023), but atherosclerosis appears earlier in men than in women. Finally, we do not know at what point in time autoimmunity to H2B develops in the atherosclerosis environment.

## RESOURCE AVAILABILITY

### Lead contact

Further information and requests for resources and reagents should be directed to and will be fulfilled by the lead contact, Changjun Yin.

### Materials availability

Materials generated in this study can be requested from the lead contact.

### Data and code availability

The BCR sequences and associated metadata for each sample, including mouse ID, tissue, genotype, mutation summary, isotype summary, and clonality for BCR repertoire sequencing, as well as the count matrix and metadata (tissue, cluster, cell type, and BCR-related information) for scRNA-seq data, and IP proteomics data have been deposited in Figshare (<u>10.6084/m9.figshare.27022876</u>). The paired CDR3 sequencing data are available from the corresponding authors on reasonable request. The public mouse GC B cellsdata used in our study can be downloaded from the Gene Expression Omnibus under accession numbers <u>GSE154634</u>. All data are available in the main text or the supplementary materials.

## ACKNOWLEDGMENTS

We thank C.E. Busse and H. Wardemann from the German Cancer Research Center, Heidelberg, Germany for their advice regarding PCR primers used for single cell BCR cloning; we thank C. Wen from the Institute of Precision Medicine, The First Affiliated Hospital of Sun Yat-sen University for illustrations; The authors thank Dr. Ignasi Forne of the Protein analysis unit at the BioMedical Center, LMU, Munich, for his help with the MS measurements; we thank Georg Wick of the University of Innsbruck for discussions related to the nature of autoimmunity in atherosclerosis.

## AUTHOR CONTRIBUTIONS

A.J.R.H. and C.Y. conceived and conceptualized the study. A.J.R.H. and C.Y. designed, performed, analyzed and supervised experiments. A.J.R.H., C.Y. and K.L. analyzed data and wrote the manuscript with input from all authors. C.Z. performed and analyzed single cell BCR cloning, antigen identification and vaccination experiments; Y.R. performed antibody transfer experiments; Z.W and X.Z. performed and analyzed scRNA-seq and scBCR-seq experiments; L.L., S.W., Y.Z., J.Z., T.S., Y.L., S.L., M.H., Z.M., M.H., X.B. performed and analyzed experiments; K.D. and D.H. contributed to methodology and helped writing; C.Z., L.L., S.L., A.I. performed and analyzed IP-MS experiments; C.Z., S.N., L.M. were involved in human carotid plaque analysis; C.Z., J.Z., J.X., J.Z., Y.W., H.H., L.H., D.S. contributed to human serum studies: S.S., C.W., S.K.M. performed and supervised experiments; Z.W. and D.S contributed to data analysis and statistics; R.J.M.B.-R. contributed to writing, review and editing the manuscript. All authors contributed to data interpretation.

## DECLARATION OF GENERATIVE AI AND AI-ASSISTED TECHNOLOGIES

During the preparation of this work, the authors did not use AI-based tools.

## SUPPLEMENTAL INFORMATION

**Document S1. Figures S1–S6 and Table S2** (this is the main PDF)

**Table S1. Immunoglobulin gene repertoire and reactivity of antibodies from WT and Apoe^-/-^ mice (table S1 provided as separated excel file).**

## FIGURE TITLES AND LEGENDS FOR SUPPLEMENTAL FIGURE

**Figure S1: ScRNA-seq/scBCR-seq pairing reveals tolerance dysfunction in ATLOs**

(A) pairing of scRNA-seq/scBCR-seq approach to compare B cell subtype transcriptomes of RLN B cell subsets of both genotypes and ATLOs.

(B) 12,854 B cells were grouped into six subsets resulting from UMAP analyses based on their top 2000 most highly expressed genes; B cell subset distribution in the UMAP plot of WT RLNs, Apoe^−/−^ RLNs and ATLOs as naїve B cells, newly activated B cells, early GC B cells, late GC B cells, memB cells and PCs;

(C) Dotplot visualization of transcript profiles of B cell subtypes; dotplots show marker signature genes to define six B cell subsets: IgM, IgD, Sell (selectin L) and early B cell transcription factor 1 (Ebf1) for naїve B cells; CD69 for activation; eukaryotic translation initiation factor 5a (Eif5a) and myelocytomatosis oncogene (Myc) for early GC B cells; activation-induced cytidine deaminase (Aicda) and proliferation-related Ki67 antigen (Ki67) for late GC B cells; CD38 and zinc finger E-box-binding homeobox 2 (Zeb2) and zinc finger and BTB domain containing 20 (Zbtb20) for memB cells; and Sdc1 (CD138) and X-box binding protein 1 (Xbp1) for PCs. 6 major B cell/PC subsets were identified across SLOs and ATLOs according to their prototypical markers, that is i. naїve B cells (IgM^+^, IgD^+^, Sell (Selectin L)^+^, Ebf1^+^); ii. newly activated B cells (Nac B cells, IgM^+^, IgD^+^, Sell^+^, CD69^+^); iii. early GC B cells (Eif5a^+^, Myc^+^, Aicda^-^, Ki67^-^); iv. late GC B cells (Eif5a^+^, Aicda^+^, Ki67^+^); v. memB cells(Zeb2^+^, Zbtb20^+^); and vi. PCs (Xbp1^+^, Sdc1^+^);

(D) UMAP visualization of clonally expanded B cell subsets; distribution of clonally expanded B cells. Clonally expanded B cells in red, not clonally expanded B cells in grey. B cells with identical IgH CDR3 amino acid sequences are considered as clonally expanded;

(E) Volcano plots show differentially expressed genes (DEGs) between two tissues. Apoe^-/-^ RLN B cells *vs* WT RLN B cells, and ATLO B cells *vs* Apoe^-/-^ RLN B cells were compared.

(F) Dotplots showed abnormal gene expression signatures in ATLO B cell subsets; gene expression profiles of 6 B cell subsets were determined as described in Methods; genes with |Log2(fold change)|>0.25 and adjusted P value<0.05 were considered as DEGs. The adjusted P value was determined by two-sided Wilcoxon-Rank Sum test with Bonferroni correction.

(G) Number of immunoglobulin heavy chain (IgH) mRNAs expression per PC; scRNA-seq/BCR-sec pairing allows to quantify IgH transcripts per PC. Statistical analyses were performed using Kruskal-Wallis H Test with Benjamini-Hochberg correction.

(H-I). ATLO PC transcripts involved in key protein secretion-regulating genes and function-related genes in Apoe^-/-^ RLNs vs ATLOs; Violin plots show PC protein secretion-regulating gene profiles and function-related gene expression in different tissues. Statistical analyses were performed using the Kruskal-Wallis H Test with Benjamini-Hochberg correction.

(J) B cell transcripts indicate tolerance dysfunction in ATLOs. See also Figure S1.

**Figure S2: Modeling of cell-cell interactions suggests aberrant regulation of ATLO B cell immunity**

(A) Aberrant interactions between myeloid-cell subtypes and B cells. Cell-cell interactions between myeloid-cell subsets and B cells were modeled as described in Methods.

(B) Aberrant interactions between T cell subsets and B cells. Interactions between T cell subsets and B cells were modeled as described in Methods. Chi-square test with Benjamini-Hochberg correction was used to perform statistical analysis for A-B.

(C-E) Aberrant interaction maps in different tissues. Predicted interactions between Trem2high macrophages (C), tolerogenic DCs (D), and T_reg_ cells (E) with naïve B cells, Nac B cells, early GC B cells, late GC B cells, memB cells, and PCs in WT RLNs, Apoe^-/-^ RLNs, and ATLOs. Color scale indicates normalized expression, dot size indicates significance (-log10 p value), x represents no predicted interactions was detected.

(F) Aberrant myeloid-cell and T cell interactions with B cell subsets indicate dysfunction of major immune checkpoints in ATLOs. Red arrows indicate aberrant interactions between myeloid cell and T cell interactions with B cell subsets.

See also Figure S2.

**Figure S3: B cell transcript signatures are dysregulated in ATLO GCs**

(A) Integration of GC B cells from WT RLNs, Apoe^-/-^ RLNs, and ATLOs with GC B cells in spleens of vaccinated WT mice; GC B cells of vaccinated WT mice (obtained from a publicly available databank: GSE154634) were integrated into our own GC B cells from unvaccinated WT GC B cells isolated from RLNs or Apoe^-/-^ RLNs or ATLOs. B cells from our own data are colored, cells from their spleen counterparts of vaccinated mice (GSE154634) were labeled in grey.

(B) Identification of 7 GC B cell subtypes in all pooled GC B cells; All GC B cells are grouped into 7 subtypes following in tSNE map analyses based on their top 2000 highly expressed genes; 7 GC B cell subtypes are further grouped into light zone (LZ) GC B cells(clusters 0, 4, 6), dark zone (DZ) GC B cells(clusters 1, 2, 3), and resting GC B cells(cluster 5).

(C) tSNE map shows cell cycle markers (G1, G2M, and S) phase of GC B cells from pools of GC B cells.

(D) Prototypic marker genes to profile LZ, DZ and resting GC B cells in pooled GC B cells; dot plot shows expression of GC B cell-related markers in 7 GC subtypes obtained from vaccinated WT mice plus GC B cells of WT RLNs, Apoe^-/-^ RLNs, and ATLOs.

(E) Prototypic marker genes expressed by GC B cells obtained from pooled of WT RLNs, Apoe^-/-^ RLNs, and ATLOs. Spleen GC B cells from GSE154634 were not included in this assay. Violin plots show representative gene expression levels associated with GC B cell subsets including Ki67 (proliferation), Aicda (enzyme mediates BCR mutation and class switching), CD83 (activation) and CXCR4 (migration) in WT RLNs, Apoe^-/-^ RLNs, and ATLOs; Each dot represents one single B cell.

(F) DEGs in GC B cells in WT RLN/Apoe^-/-^ and Apoe^-/-^ RLN/ATLO comparisons; Volcano plot of DEGs of WT RLN GCs, Apoe^-/-^ RLN GCs, ATLO GCs. Genes with |Log2(fold change)|>0.25 and adjusted P value<0.05 were considered as DEGs. The adjusted Pvalue was determined by two-sided Wilcoxon-Rank Sum test with Bonferroni correction.

See also Figure S3.

**Figure S4: ATLO GC B cells show increases in GC B cell-associated cytokine pathways and defects of tolerance maintenance**

(A) Schematic of experimental approach to identify differentially regulated genes in SLO vs ATLO GC B cell subsets; identification of DEGs of GC B cells was followed by gene set enrichment analyses (GSEA).

(B) Normalized enrichment scores (NES) of GO terms in ATLO GC B cells vs their counterparts in WT RLNs.

(C-D GSEA assays of selected GO terms enriched in ATLO LZ GC B cells and ATLO GC B cells vs WT and Apoe^-/-^ RLNs. The x-axis shows genes (vertical black lines) represented in the GO terms and the color indicated positive (red) or negative (blue) correlation. The green line connects gene and enrichment score of selected biological pathways. NES: normalized enrichment score.

(E) Dotplot shows individual gene expression profiles involved in different biological functions for each GC B cell subset.

See also Figure S4.

**Figure S5: H2B autoimmune pathway does not impact blood lipids.**

(A) Experimental design of the H2B vaccination mouse model; vaccination using recombinant H2B in young Apoe^-/-^ mice. Apoe^-/-^ mice were vaccinated with recombinant H2B with adjuvant (termed as H2B vaccine), adjuvant alone, or PBS alone at the age of 8 weeks with a boost at the age of 12 weeks. The experiment was completed at 32 weeks. Blood was collected at multiple time points before and after the injection; aortic root sections were stained for Oil red O (ORO). Plaque sizes were quantified as described in Methods; Scale bar 500 μm. PBS control (n = 7 mice), adjuvant control (n = 7), H2B + adjuvant (n = 8).

(B-C) Vaccination of H2B in hyperlipidemic mice does not affect blood lipids (B) or body weight (C); blood was collected at the end of the experiment; body weight was measured at multiple time points before and after the injection.

(D-E) A6 antibody transfer does not affect blood lipids and body weight in Apoe^-/-^ few with chow diet or high fat diet. Experimental design of the A6 transfer to mice. Apoe^-/-^ mice fed with chow diet (D) or high fat diet

(E) were used. Triglyceride (TG), cholesterol (CHO), high-density lipoprotein cholesterol (HDL), and low-density lipoprotein cholesterol (LDL) were measured.

See also Figure S5.

**Figure S6 Hypothetical choreography of the B cell tolerance breakdown in ATLOs**

A fraction of naïve B cells after entering the ATLO find cognate (auto)antigens and - upon subsequent help received from CD4^+^ T helper cells and TFH cells - become newly activated B cells followed by proliferation; some of these B cells may differentiate into low-affinity memB cells or low-affinity PCs without entering the GC; others are recruited into the GC to advance to low-affinity early GC B cells; low-affinity early GC B cells subsequently engage in the GC high-affinity maturation cycle via interaction with cognate (auto)antigens bound to the surface of FDCs and they also receive helper signals from TFHs; after repeated cell cycles, some of these GC B cells become high-affinity memB cells or high-affinity PCs (while others are deleted or silenced via tolerance mechanisms; after the GC reaction, high affinity PCs form survival niches while others home to the bone marrow and SLOs; similarly, high-affinity memB cells emigrate to home into peripheral tissues and are ready to protect the organism to mount a vigorous antibody response upon a second exposure to antigen. Multiple tolerance checkpoints are compromised at ATLOs: **①** Exaggerated pre-GC myeloid/B cell and T cell/B cell interaction; **②** exaggerated B cell recruitment of newly activated B cells into the GC and less differentiation into low-affinity PCs; **③** loss of GC negative selection tolerance allow formation of high-affinity autoantibodies; **④** abnormally large PC niches. Tissue-specific tolerance dysfunction is consistent with the possibility that ATLOs are preferred lymphoid organs to generate atherosclerosis-directed high-affinity autoantibodies directed against antigenic self-molecules. See also Figure S6.

## STAR★METHODS

### KEY RESOURCES TABLE

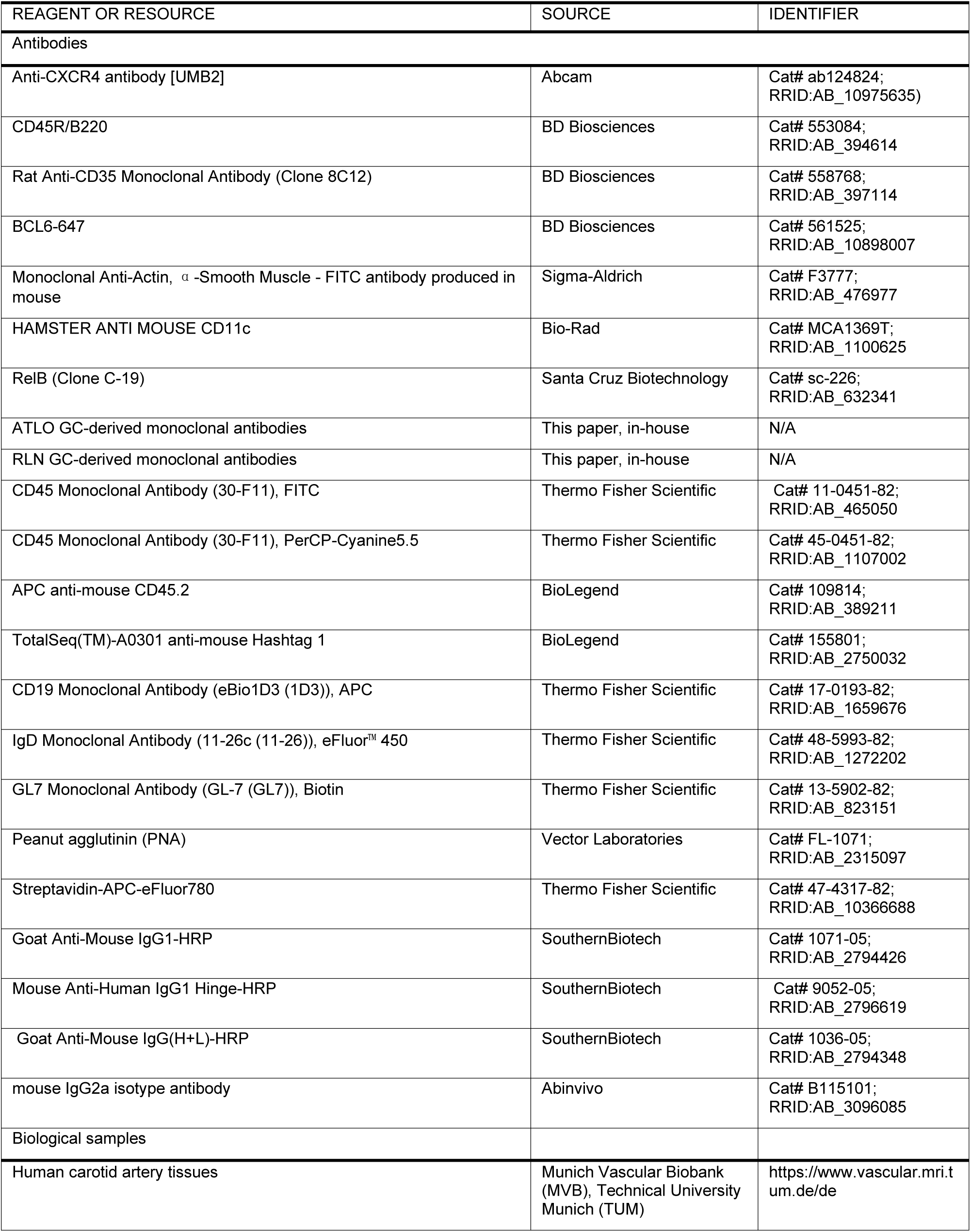
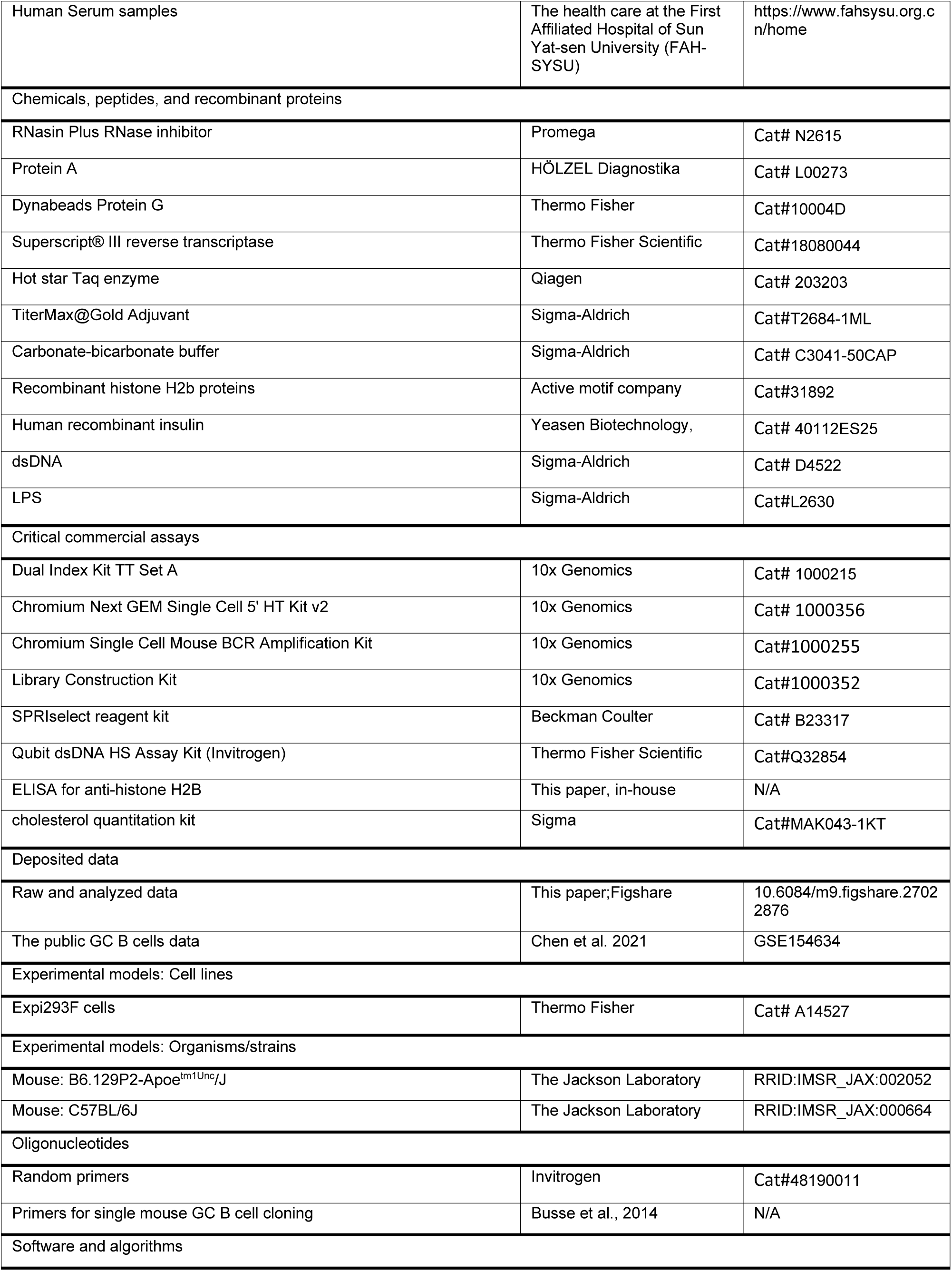
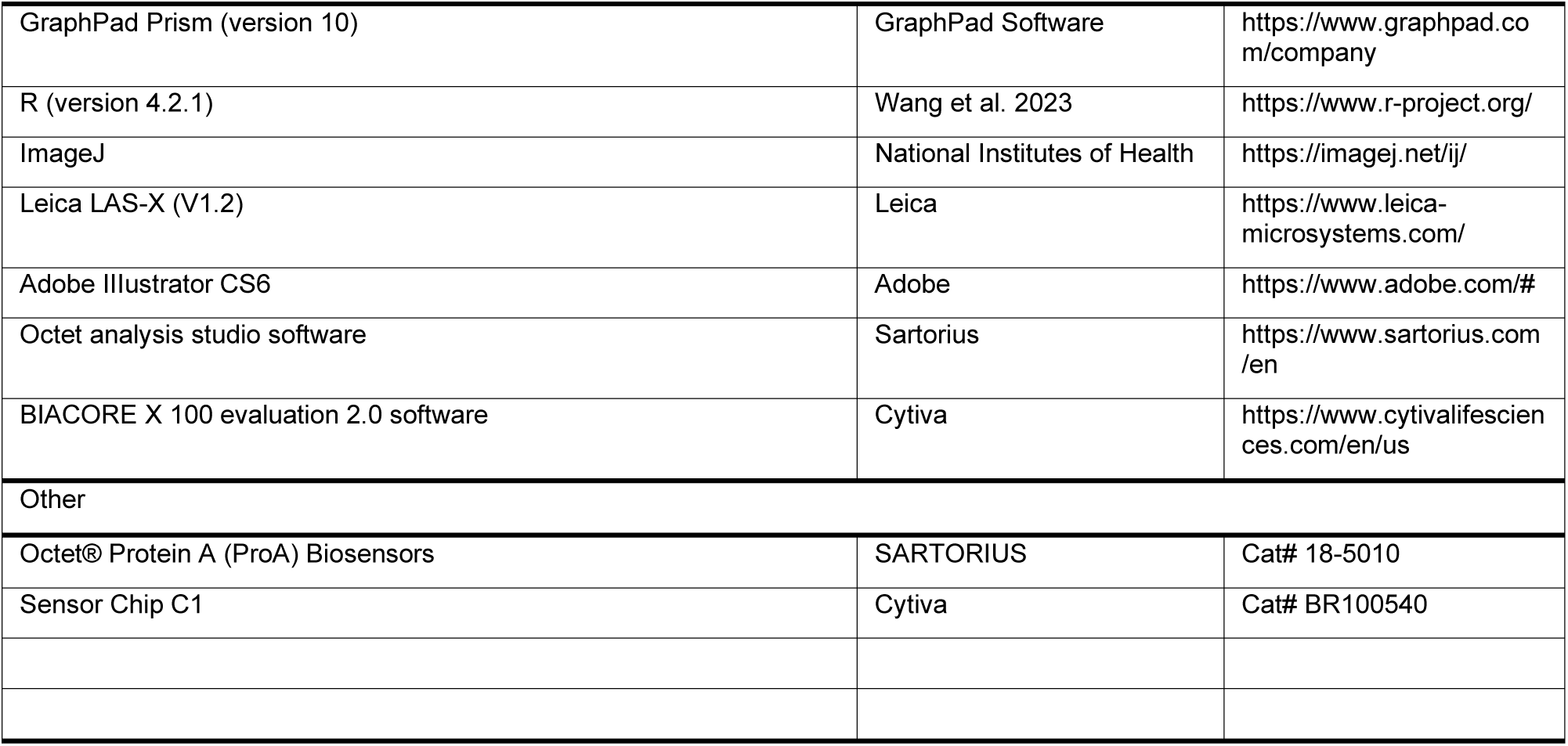

### EXPERIMENTAL MODEL AND STUDY PARTICIPANT DETAILS

#### METHOD DETAILS

##### Materials and Methods

###### Mice

WT and Apoe^-/-^ mice on a C57BL/6 background were raised in a specific pathogen-free environment at the animal facilities of Ludwig-Maximilians-Universität München (LMU) until 78 weeks of age or at the animal facilities of the First-affiliated Hospital of Sun Yat-sen University at 8 - 32 weeks of age. Mice were given a standard rodent chow or a high fat diet containing 40% fat, 1.25% cholesterol, and 0.5% sodium cholate (D12109C, Research Diets, New Brunswick, NJ, USA) and maintained in a climate-controlled room at 23 °C and 60% relative humidity, with a 12-hour light/dark cycle. The animal procedures were authorized by the Regierung of Oberbayern or by the First-affiliated Hospital of Sun Yat-sen University, following the guidelines of the local Animal Use and Care Committee and the National Animal Welfare Laws.

###### Tissue collection

Mice were anesthetized using hydrochloride and xylazine hydro-chloride mixed solution (2:1) through intraperitoneal injection. To collect tissues for BCR repertoire mapping: the peritoneal cavity was washed with 10 ml of FACS buffer (DPBS buffer with 2% foetal bovine serum) to obtain lavage, while blood was collected by cardiac puncture. Following perfusion using 10 ml EDTA buffer (5 mM), 20 ml PBS and 20 ml FACS buffer, RLNs, spleen and ATLO were isolated as described previously(Hu et al., 2015a; Wang et al., 2022). Bone marrow was collected by flushing the femur and tibia with 10 ml of FACS buffer. All harvested tissues were immediately placed on ice for subsequent processing. ATLO tissues from the diseased aorta for single cell RNA sequencing and single cell BCR sequencing were collected as described previously(Wang et al., 2023): In short, adipose tissue and paraaortic lymph nodes were carefully removed. The whole aorta was dissected and collected in cell culture dishes with pre-cooled FACS buffer. The aorta was opened in the longitudinal direction, and plaques were carefully removed using curved forceps (Dumont 5/45, Fine Science Tools) under the dissecting microscope. The remaining aorta was collected as ATLOs.

Aortas with no ATLOs (as indicated with no atherosclerosis or few atherosclerotic plaques in the abdominal aorta) were excluded from the subsequent studies. Two separate cohorts of mice were used: (1) pools of three Apoe^-/-^ and three WT mice were collected for ATLOs and RLNs, and pools of five Apoe^-/-^ and five WT mice were collected for blood in the first cohort; (2) cell hashtags (hashtags, BioLegend, TotalSeq C) were used to label four Apoe^-/-^ and four WT mice individually to collect ATLOs and RLNs as reported previously(Wang et al., 2023).

###### Human carotid artery tissues

Human carotid atherosclerotic plaques were obtained from patients with high-grade carotid artery stenosis (> 70%) after carotid endarterectomy, and were provided by the biobank Munich, Technical University Munich (TUM) according to the guidelines as described previously(Yin et al., 2019). The study was performed according to the Guidelines of the World Medical Association Declaration of Helsinki. The ethics committee of Faculty of Medicine, Technical University of Munich (TUM) approved the study, and written informed consent for permission to be included into the Munich Vascular Biobank was given by all patients.

###### Clinical cohort methods

We performed a cross-sectional study of 495 individuals of the general population who undergoing routine medical screening of preventive health care at the First Affiliated Hospital of Sun Yat-sen University (FAH-SYSU), China. The study was approved by the Clinical Research and Laboratory Animal Ethics Committee, The First Affiliated Hospital of Sun Yat-sen University, and all participants provided their informed consent for participation. The baseline demographic and clinical characteristics of the participants are summarized in **Supplementary Table S2**. Exclusion criteria included patients with previously diagnosed cardiovascular diseases, strokes, autoimmune diseases requiring systemic treatment, active malignancies, and severe kidney or liver dysfunction. Blood samples were collected to assess general clinical laboratory parameters and serum titers of anti-H2B antibodies. Additionally, as part of medical screening program, chest 3-dimentional computed tomography (CT) scans were performed voluntarily for general diagnostic purposes. Non-contrast enhanced chest CT scans were performed as part of the clinical assessment with a 64-row multidetector CT scanner. The scanned area started from the apex to the base of the lungs, including the heart and aortic arch, and was acquired in the craniocaudal direction during inspiration. The scan parameters were as follows: tube voltage, 120 kV and tube current, 200 mAs; beam pitch, 0.828; rotation time, 0.5 seconds; beam collimation, 64 mm × 0.5 mm. Thoracic aortic wall calcification (TAC) was considered a binary outcome defined by the presence of hyperattenuating signals (above the threshold of 100 Hounsfield units and assessed by experienced radiologists) in an area of at least three adjacent pixels within the boundaries of the thoracic aorta (ascending, arch, and descending aorta). Valid data for CT scan evaluation were available for 418 individuals. The univariate association between anti-H2B antibody titers and TAC was analyzed by univariate logistic regression and analysis of the receiver operating characteristic (ROC) curves under the assumption of non-parametric distribution. For multivariate analysis, we used binary logistic regression models. The model included log-transformed anti-H2B antibody titer (to approximate Gaussian distribution), sex, age, body mass index (BMI), systolic blood pressure, diastolic blood pressure, HDL cholesterol, LDL cholesterol, triglycerides, estimated glomerular filtration rate (eGFR), homocysteine, and hemoglobin A1c (HbA1c). Total cholesterol levels were excluded due to significant collinearity with LDL cholesterol. The results of the simultaneous-entry model were confirmed using a backward logistic regression approach. In this model, sex, age, anti-H2B antibody titer, LDL cholesterol, and HbA1c were retained in the final step as independent predictors.

###### Single cell suspension preparation

Tissues were dissected into small pieces and mashed to allow passage through 70 µm^2^ strainers. Aorta samples were subjected to digestion with 2.5 ml enzyme cocktail (400 U ml^−1^ collagenase type I, 10 U ml^−1^ collagenase type XI, 60 U ml^−1^ hyaluronidase, 60 U ml−1 DNase I and 20 mM HEPES in DPBS) at 37 °C for 45 min with slow shaking prior to filtering. The filtered cell suspension was subsequently centrifuged at 300G for 5 min. Following the removal of the supernatant, 5 ml Ack lysis buffer (0.15 mM NH_4_Cl, 1 mM KHCO_3_, 0.1 mM Na_2_EDTA, pH 7.2-7.4) was added to spleen, blood, and BM samples to induce lysis of red blood cells. This lysis process lasted for 5 min at room temperature (RT), followed by two rounds of washing to eliminate cell debris and residual lysis buffer. Cells were then resuspended in 1 ml pre-cooled FACS buffer and passed through 100 µm^2^ filters before cell counting. The cell number was determined using a Neubauer chamber under the microscope or by an automated cell counter (TC20, Bio-Rad).

###### Single cell isolation for single-cell profiling

Samples were prepared as reported previously(Wang et al., 2023): Following the single cell suspension preparation, cells of each tissue were stained with fixable viability dye (1:1,000 dilution, eBioscience) in PBS at 4 °C for 20 min. After washing and blocking steps, cells were stained with CD45 labeled in fluorescence (1:200 dilution, BD Bioscience, clone 30-F11, for pooling cohort) or CD45.2 (1:200 dilution, BioLegend, clone 104, for hashtag cohort) with or without hashtagged CD45/MHCI antibody (1:50 dilution, BioLegend, TotalSeq C) at 4 °C for 30 min. Single CD45^+^ live cells were sorted using a flow cytometer (FACSAria III, BD Biosciences). Single CD45^+^ live cells were sorted into a cold tube pre-rinsed with FACS buffer using a flow cytometer (FACSAria III, BD Biosciences). Cell number and viability were determined by using of automated cell counter (TC20, Bio-Rad). The cells were maintained on ice until loaded to the Chromium controller.

###### Single GC B cell isolation for PCR cloning

Following preparation of single cell suspensions from ATLOs and RLNs, cells were stained with FVD (APC-Cy7-conjugated, 1:1000 dilution, eBioscience) to label live cells at 4 °C for 20 min), followed with CD45 PercP-Cy5.5 (30-F11, eBioscience), CD19 APC (1D3, eBioscience), IgD-eFluor 450 (11-26c, eBioscience), GL7-Biotin (GL7, eBioscience), PNA-FITC (Vector Lab), Streptavidin-APC-eFluor780 (eBioscience) in pre-cooled FACS buffer. Single GC B cells (FVD^-^CD45^+^CD19^+^IgD^-^GL7^+^PNA^+^) were sorted into pre-cooled 96-well plate containing 4 µl lysis buffer (final concentrations: 0.5 x DPBS without Ca/Mg, 3 U/µl RNasin Plus RNase inhibitor (Promega, Cat. no. N2615), 10 mM DTT) as described previously(Srikakulapu et al., 2016). Then, 96-well plates were sealed with Microseal ‘F’ Film (BioRad) and immediately frozen on dry ice before storage at −80 °C.

###### Single B cell cDNA synthesis for PCR cloning

Total RNA from a single GC B cell on 96-well plate was reverse transcribed in RNA/DNA free water using 128 ng random primers (Invitrogen 48190-011), 16.2U RNasin Plus RNase inhibitor (Promega), 0.3% IGEPAL CA-630 (Sigma-Aldrich), 100mM DTT, 12.6 mM dNTP, and 50 U Superscript® III reverse transcriptase (Invitrogen) as described previously(Srikakulapu et al., 2016). Reverse transcription reactions were performed at 42 °C for 5 min, 25 °C for 10 min, 50 °C for 60 min and 94 °C for 5 min. cDNA was stored at −20 °C before PCR.

###### Cloning of paired heavy and light chains of BCR from single GC B cells

PCR primers used for single mouse GC B cell cloning were synthesized by Metabion company (Munich); the detailed sequences of primers were described previously(Busse et al., 2014; Tiller et al., 2009; von Boehmer et al., 2016). The final concentration of all primers was 100 μM. Mouse BCR heavy chain and BCR light chains were amplified independently by nested PCR starting from 4 µl cDNA as template. All PCR reactions were performed in 96-well plates in a total volume of 42 µl PCR buffer containing 0.3 µM/µl dNTP, 0.3 µM each primer or primer mix, and 0.05 U/µl Hot star Taq enzyme (Qiagen, Cat. no. 203203). Semi-nested or nested second round PCR was performed with 4 μl of unpurified first round PCR product at 95 °C for 15 min followed by 50 cycles of 94 °C for 30 s, 60 °C (Igh) or 58 °C(Igl) for 30 s, 72 °C for 45 s, and final incubation at 72 °C for 10 min. PCR products were purified by purification kit (Qiagen) according to the manufacturer’s instructions. Purified DNA were sequenced by Sanger sequencing at the Faculty of Biology, LMU (http://www.gi.bio.lmu.de/sequencing), Munich. BCR sequences were further compared to the germline line BCR sequences from IMGT databank (http://www.imgt.org/IMGT_vquest/vquest?livret=0&Option=mouseIg). Paired BCR heavy chain and light chain sequences were used to generate monoclonal antibodies.

###### Single-cell profiling library preparation for gene expression and B cell receptor (BCR) sequencing

Cell concentrations were adjusted to 700-1,200 cells per microliter according to the protocol of 10X Genomics 5′ library & gel bead kit. Single-cell suspensions were mixed with nuclease-free water and 5′ single-cell RNA master mixture, then loaded into a Chromium chip with barcoded gel beads and partitioning oil. The chip was placed in the Chromium controller or Chromium X to generate gel beads in emulsion (GEMs). cDNA was obtained from 100–180 μl GEMs/sample by reverse-transcription reactions: 53 °C for 45 min, 85 °C for 5 min, then maintained at 4 °C. cDNA products were purified and cleaned using Dynabeads. cDNA was amplified by PCR: 98 °C for 45 s; 98 °C for 20 s, 67 °C for 30 s, 72 °C for 1 min and amplified for 13–16 cycles; then, 72 °C for 1 min. Amplified PCR products were cleaned and purified using SPRIselect reagent kit (B23317, Beckman Coulter). The concentration of cDNA library was determined by Qubit dsDNA HS Assay Kit (Invitrogen). The single-cell transcriptome library and B cell V(D)J library were prepared using Chromium single cell V(D)J reagent kits following the standard guidance. 3000 single cells were expected be recovered for each sample. The reverse transcription and following library construction were performed using C1000 Touch thermal cycler (BIO-RAD). Totally 14 cycles were set for cDNA amplification. The concentration of each library was determined using Qubit 3.0 fluorometer and NEBNext Library Quant kit, and the size range of final libraries was checked on a fragment analyzer (Advanced Analytical Technologies). 5’ gene expression libraries or BCR libraries were pooled together with the same quality for sequencing.

###### Next-generation sequencing

The concentration of pooled libraries was quantified using both Qubit 3.0 fluorometer and NEBNext Library Quant kit. The size of libraries was assessed using the Bioanalyzer (Agilent). Only libraries of high quality were loaded onto the Illumina platform. The pooled libraries of BCR repertoire mapping, samples with 30% PHiX control library spike-in (Illumina) were loaded onto MiSeq platform (Illumina) and subjected to 2 x 300 bp paired-end sequencing at IMGM company (Munich, Germany; https://www.imgm.com/). For the first cohort of pooled samples of scRNA-seq and scBCR-seq, 5′ gene expression libraries and BCR libraries were sequenced separately at IMGM. Around 1% PHiX control library spike-in (Illumina) was added to 5′ gene expression libraries and sequenced by a NextSeq high-output reagent kit v.2.5 (150 cycles) using the Nextseq 500 platform (Illumina). Sequencing was performed as an asymmetrical manner: read 1 was adjusted to 26 bp, read 2 to 98 bp and the index 1 to 8 bp. BCR libraries together with 1% PHiX control library spike-in were sequenced in a symmetrical manner (150 bp + 150 bp, paired end). For the second cohort of hashtagged samples, 5′ gene expression libraries, BCR libraries and cell hashtag libraries were sequenced together using the NovaSeq 6000 platform at IMGM (https://www.imgm.com/). Three types of libraries were pooled at the ratio of 40:10:1. Additionally, 1% of PHiX library (Illumina) was spike-in as control. Sequencing was performed in an asymmetrical manner: read 1 was adjusted to 28 bp plus 10 bp for i7 index, 10 bp for i5 index, and read 2 to 90 bp.

###### Single-cell sequencing data pre-processing analysis

The NGS sequencing data were de-multiplexed based on Illumina index using 10x Genomics Cell Range mkfastq pipeline (version 3.0.0). The 5’ gene expression matrix was generated by aligning the data to mm10 genome reference using STAR aligner built-in Cell Ranger. Count matrix of each sample was loaded into R (version 4.2.1) software using the *Seurat* (version 3.4.0) package(Satija et al., 2015). Cells contained fewer than 200 genes, higher than 4,000 detected genes or > 8% of mitochondrial genes were filtered out as suspected doublets or debris. Data from different tissues were log-normalized, and 2,000 highly variable features were calculated to integrate data. After integration, principal component analysis (PCA) was performed for following dimensional reduction and clustering. 20 dims were included to reduce the dimension and assign clusters. The clustering result was visualized using uniform manifold approximation and projection (UMAP) dimensionality reduction methods built in *Seurat*. In order to focus on B subsets, clusters highly expressed *Cd19* or *Sdc1* (CD138) genes were extracted and re-clustered for the following analyses. GC B cells were extracted and integrated with published data (GSE154634) following the integration pipelines provided by the *Seurat* package(Chen et al., 2021). 0.2 resolution was introduced to nonsupervisory cluster all integrated GC B cells. The phase of cell cycle was annotated by calculating the expression of cell cycle-related markers provided by Tirosh et al. using the cell cycle scoring pipelines in the *Seurat* package(Kowalczyk et al., 2015). PCs were extracted to be re-clustered base on 0.6 resolution. The BCR sequences were assembled and annotated referring to GRCm38 reference. The information of BCR sequences in each B cell was integrated into metadata of 5’ gene expression data for further analysis.

###### Differentially expressed genes (DEGs) and pathway enrichment analyses

In scRNA-seq analysis, DEGs for each tissue or each B cell subset were determined with the following criteria: |Log(fold change, FC)| > 0.25 and adjusted p-value < 0.05. The normalized expression of DEGs in each cell was visualized with DoHeatmap function built in the *Seurat* package. DEGs obtained from each B cell subsets were parsed to the *clusterProfiler* package for functional enrichment and multiple comparisons. Terms showing a false discovery rate (FDR) less than 0.05 were considered as significant. Selected enriched terms were illustrated by *ggplot2*. The results of gene set enrichment analysis (GSEA) were also obtained using the *clusterProfiler* package and the GSEA plot for selective GO term was visualized by the *enrichplot* package.

###### Cell-cell interaction analyses

The count of cell-cell interactions was predicted and visualized with *CellChat* (version 2.1.2) package in R software. The ligand-receptor interaction pairs were analyzed using *CellPhoneDB* (version 5). Specifically, gene symbols in our dataset were converted to corresponding human gene symbols using *biomaRt* (version 2.54.1). Genes that did not match human genes were removed, and in cases where multiple human genes matched, only one was retained. The gene expression count matrix and cell annotation data (metadata) were used as input for analysis in each tissue. The cell-cell interaction analysis was performed using Python (3.8.13) with default settings. For visualization, interaction pairs with an interaction score greater than 0 and p values less than 0.05 were considered significant. P values less than 0.0001 were represented as 0.0001 in the dot plot for better visualization.

###### Immunization Apoe^-/-^ mice with Histone 2B protein

Equal amounts of TiterMax@Gold Adjuvant (Sigma, T2684-1ML) and recombinant Histone H2B protein (Active motif company, https://www.activemotif.com/) was used to prepare the adjuvant – H2B emulsion. After preparing the adjuvant – H2B emulsion, 8-week-old Apoe^-/-^ mouse either received 200 μl PBS alone, adjuvant, or adjuvant – H2B (25 µg/mouse) emulsion by intraperitoneal injection. Four weeks after the primary injection, H2B (25 µg/mouse) in 200 μl PBS was used as the booster injection. The control group received the same volume of PBS. Every 1 or 2 weeks, the body weight was measured. The blood was collected through the tail vein to measure the titer of antibodies and cholesterol levels. Animals were sacrificed at 32 weeks of age. Whole aortas and aortic roots were collected to quantify atherosclerosis.

###### Transfer of A6 antibody in Apoe^-/-^ mice

Plasmids encoding the heavy (IgH) and light (IgL) chains of the A6 antibody were transfected into Expi293F cells (Thermo Fisher, A14527) using polyethyleneimine (PEI, Sigma-Aldrich) as the transfection reagent. Seven days post-transfection, the cell culture supernatant was harvested and purified using a Protein A Sepharose (Sigma, P3391) chromatography column. The eluate was collected and transferred into pre-soaked dialysis membranes (Spectrum Labs, 132554), which were placed in PBS and gently stirred overnight at 4 °C. The dialyzed antibodies were then concentrated using Amicon Ultra Centrifugal Filters with a 30 kDa molecular weight cutoff (Millipore, UFC903024). IgG concentrations were determined by ELISA, and the purity of IgG was assessed by SDS-PAGE (4%–12%). Low/no endotoxin levels (< 5 EU/ml) were confirmed by endotoxin assay kit (Thermofisher, A39552). To study the effect of A6 antibody in atherosclerosis mouse models, including Apoe^-/-^ mice fed with chow diet and Apoe^-/-^ mice fed with high fat diet. 12-week-old Apoe^-/-^ male mice were maintained on a chow diet for 20 weeks. Every 10 days, 100 μg A6 antibody (mA6-IgG2a) or mouse IgG2a isotype antibody (Abinvivo, B115101) or equal volume of PBS control were intravenously (i.v.) administered. After 20 weeks of treatment, the mice were killed for tissue collection. To study the effect of A6 in a high fat diet-induced atherosclerosis mouse model. 10-week-old Apoe^-/-^ male mice were fed high fat diet for 6 weeks. 100 μg A6 antibody (mA6-IgG2a) or mouse IgG2a isotype antibody (Abinvivo, B115101) or equal volume of PBS control were intravenously (i.v.) administered at 12-week. After 4 weeks of treatment, the mice were killed for tissue collection.

###### Atherosclerosis quantification

En-face Sudan IV and Oil-red-O (ORO) staining were used to analyse the mouse aortas as reported previously(Yin et al., 2019). To assess lipid-rich areas in the whole aorta (*en face* staining), arch, thoracic, and abdominal parts, a digital camera (Leica) was used to capture images with a standard bar. Oil-red-O (ORO) staining was used to analyse atherosclerotic plaque areas. The Thunder imaging system (Leica) was used to capture ORO staining images. Image J was used to measure the percentages of lipid deposition in the whole aorta, arch, thoracic, and abdominal parts and quantified atherosclerotic lesion areas as previously(Yin et al., 2019).

###### Determination of antibody-antigen binding affinity by Surface Plasmon Resonance (SPR)

The surface plasmon resonance analyses were performed using BIAcore X 100 (Cytiva, Freiburg, Germany) equipped with a research-grade C1 sensor chip. The ligand was immobilized using amine-coupling chemistry at a density of 492 resonance U on flow cell 2; flow cell 1 was left empty to be used as a reference surface. Analytes in running buffer (PBST: 0.1 % tween 20 with 0.05 % BSA in PBS, pH 7.4) were injected at a flow rate of 90 μl/min. The complex was allowed to associate and dissociate for 60 and 240 s, respectively. The surfaces were regenerated with 2 pulses (60 s) of 30 mM NaOH, 2 M NaCl, 0.3 % TritonX - 100. Responses from analyte injections (colored curves) were overlaid with the fit of the 1: 1 interaction model (shown as red curve) determined using BIACORE X 100 evaluation 2.0 software.

###### Examination of antibody-antigen binding affinity by Biolayer Interferometry (BLI)

The bio-layer interferometry analyses were performed using an OCTET R8 (SARTORIUS, Otto-Brenner-Str. 2037079 Göttingen, Germany) equipped with research-grade Protein A Biosensors (18-5010, SARTORIUS). Different antibodies (e.g., A6, isotypes) were coupled to Protein A Biosensors, and antigen-antibody affinity was analyzed using the same concentration gradient of histone 2B as the analyte. Protein A biosensors were hydrated for 10 mins in DPBS (70011044, Gibco) before loading 10 µg/ml antibody for 120 s. After a 60 s baseline step, protein analyte was associated for 180 s to the sensor followed by a 180 s dissociation step. KD values were determined with a global fit 1:1 binding algorithm using the Octet analysis studio software.

###### Western blot

Aorta were lysed in ice-cold radioimmunoprecipitation assay lysis buffer containing a protease inhibitor cocktail after removing blood. Equal amounts of protein lysates were separated on the gel and transferred from the gel to the membrane. Membranes were subsequently blocked for 2 hr at RT in 5% BSA blocking buffer. After that, the membrane was incubated with primary antibodies. The membrane was washed with TBST and incubated with the horseradish peroxidase (HRP) conjugated secondary antibody in blocking buffer at RT for 1 hr. The last step was to wash the membrane 3 times with TBST for 10 min. The HRP chemiluminescent substrate (Thermo Fisher) working solution was prepared by mixture of equal parts stable peroxide solution and enhanced substrate solution in a clear tube (600 μl per membrane) and added to the membrane. CCD camera imaging devices (LAS3000 Imaging System, Fuji) were used to detect chemiluminescence.

###### Antibody polyreactivity assay by ELISA

Antibody binding capacity to three structural unrelated proteins were determined by ELISA, including double-stranded DNA (dsDNA), lipopolysaccharides (LPS), and insulin(Guthmiller et al., 2020). ELISA plates (Costar, 9018) were prepared by coating with 5 μg/ml of human recombinant insulin (Yeasen Biotechnology, 40112ES25) or 10 μg/ml of dsDNA (Sigma-Aldrich, D4522) or 10 μg/ml LPS (Sigma-Aldrich, L2630) in a 0.05 M Carbonate-Bicarbonate Buffer (Sigma-Aldrich, C3041-50CAP), and incubated overnight at room temperature. Subsequent blocking was achieved using 2% bovine serum albumin (BSA). Antibodies were applied at a concentration of 4 μg/ml, followed by three serial dilutions (1:4), and incubated at room temperature for one hour. Detection of bound antibodies was performed using a rabbit anti-human IgG HRP-conjugated antibody (Abcam, ab6759) and an HRP substrate (Sigma-Aldrich, T4444). Optical density (OD) at 450 nm was measured using a Multiskan SkyHigh (Thermo Fisher Scientific, USA). RA32 antibody was cloned from Apoe^-/-^ RLN GC B cells, RW49 antibody was cloned from WT RLN GC B cells, AT5 antibody was cloned from ATLO GC B cells. RA32 antibody recognized all three antigens strongly was used as high positive control. AT5 antibody did not bind to all three antigens was used as negative control. The detail aminol acid (AA) sequences of the paired full length of variable region of IgH and IgL are listed below: RA32-IgH-QVQLRQPGIELAKSGTSVKLSCKASGFTFTSYWMHWVRQRPGQGLEWIGHINPNKDGINFNERFKTKATLTVDRSSATAFMQLSSLTSADSAVYYCARWGPGYFDYWGQGTTLTVSS; RA32-IgL-DLVLTQSPASVAVSPGQRATISCRVSESIDNFGISFMNWFQQKPGQPPKLLIYFASKQGSGVPARFTGSGSGTNFSLNIHPMEEDDTAMYFCQQSKEVPYTFGGGTKLEIK; RW49-IgH-EVQLVESGGGLVKPGGSLKLSCAASGFTFSDYGMHWVRQAPEKGLEWVSYISSGSSTIYYADTVKGRFTISRDNAKNTLFLQMTSLRSEDTAMYYCARPGYYGLYAMDYWGQGTSVTVSS; RW49-IgL-DIKMTQSPSSMYASLGERVTITCKASQDINSYLSWFQQKPGRSPKTLIFRANRLVDGVPSRFSGSGSGQDYSLTISSLEYEDMGIYYCLQYVEFPHTFGGGTKLEIK;

###### Immunoprecipitation (IP) and protein mass spectrometry (MS)

Firstly, Protein A (HÖLZEL Diagnostika, L00273) and Dynabeads™ Protein G (Thermo Fisher, 10004D) were used to capture ATLO-derived antibodies through binding the Fc regions of Igs to protein A/G. 50 μl of total protein A/G beads were taken (i.e., 25 μl of protein A + 25 μl of protein G) into a clean tube and washed three times with 500 μl IP buffer (10 mM Hepes pH 7.6, 150 mM NaCl, 12.5 mM MgCl2, 0.1 mM EDTA, 0.5 mM EGTA, 10% Glycerol) for 5 min at RT. The tube was placed on the magnet and the supernatant was removed in each wash. After that, 8 μg antibodies in 500 μl IP buffer were added to couple to the 50 μl beads and incubated on a rotating wheel overnight at 4 °C. The second day, after washing 3 times with IP buffer, protein extract was added. The samples were incubated for 4 hrs on a rotating wheel at 4 °C. Then, samples were washed 3 times with IP buffer with 5 - 10 min incubations on a rotating wheel at 4 °C. The tube was added 50 mM NH4HCO3 buffer to resuspend the samples and placed on the magnet to remove the supernatant. After the second NH4HCO3 wash, 1 ml NH4HCO3 was added to re-suspend the beads and transferred to a fresh tube. The tube was placed on the magnet to remove as much supernatant as possible with 200 μl pipette tips. The samples were digested 30 minutes at 25 °C at 800 rpm with 100 μl (or enough to cover the beads) of 10 ng/μl trypsin in 1 M urea in 50 mM NH4CO3. The supernatant was collected into a fresh tube. The beads were washed twice with 50 μl of 50 mM NH4HCO3 and pooled the supernatants into the corresponding tube. DTT was added to the samples to a final concentration 1 mM DTT and digested overnight at 25 °C at 800 rpm. Following the digestion, alkylation-reduction was performed by adding 10 μl of iodoacetamide (5 mg/ml) and incubating for 30 min in the dark at 25 °C followed by 1 μl of 1 M DTT for 10 min at 25 °C. The samples were further acidified using 2.5 μl TFA. Peptides were then loaded to C18 stagetips (for desalting and purification) that were prior washed using 20 μl each of Methanol, 80% acetonitrile - 0.1 % trifluoroacetic acid (TFA), and 0.1 % TFA only. After binding of the peptides to the C18 stage tips, they were washed 3 times with 20 μl 0.1% TFA followed by a fast spin at 11000 g for 5 min. Samples were eluted by 3X 15 μl 80% acetonitrile - 0.25% TFA. They were then dried using a SpeedVac centrifuge and resuspended in 15 μl of 0.1% TFA. The desalted peptides were injected and separated on an Ultimate 3000 RSLCnano (Dionex, Germering, Germany) at Biomedical Center Munich Molecular Biology, LMU. Proteins were identified and quantified using MaxQuant v1.6.17.0. Proteins with a log2 fold change greater than 1 (indicating enrichment of the candidates in aorta sample when compared to control) were identified for each sample. Venn diagram analysis was performed to identify shared candidates by three independent aortas.

###### ELISA for anti-histone H2B in the circulation

In order to determine the titer of anti-histone H2b antibodies in the circulation of humans and mice, 100 μl purified recombinant histone H2b proteins (Active motif company, Cat: 31892) at 2 μg/ml concentration in the carbonate-bicarbonate buffer (Sigma, C3041-50CAP) were immobilized on high binding capacity microliter plates (Costar, 9018) overnight at 4 °C. Then, the microliter plates were blocked with 300 μl per well of 2 % BSA for at least 2 hr at RT. Subsequently, H2b-coated plates were incubated for 1 hr at RT with 100 μl recombinant anti-histone H2b antibody as standard (recombinant anti-histone H2b antibody was diluted stepwise to 250 ng/ml, 100 ng/ml, 50 ng/ml, 25 ng/ml, 10 ng/ml, and 5 ng/ml in antibody dilution buffer) or serum from 8, 32, 78 weeks WT and Apoe^-/-^ mice (at 1: 250 dilution) and humans with atherosclerosis (at 1: 200 dilution). The plates were incubated with HRP-conjugated secondary antibodies (1: 2000, Southern Biotech) for 1 hr at RT in the dark. In the end, 100 μl per well of TMB (Kementec Diagnostics, DY999) was used to incubate plates for 5-15 min at RT. The 2 N sulfuric acid (100 μl per well) was used to stop the ELISA reaction at an OD value between 1-3. The absorbance was measured at 450 nm using a multi-well plate reader (SpectraFluor Plus, Tecan).

The standard curve was drawn based on known recombinant anti-histone H2b antibody concentrations plotted against the absorbance values after subtracting the OD value from the background value (no antibody background). Sigmoidal 4-parameter was used to fit the standard curve in GraphPad Prism 6. The serum samples background-subtracted OD values were inputted into Prism and the concentration of serum samples were reported as ng/ml.

###### Immunofluorescence staining

For Immunofluorescence staining, tissues were dissected and embedded in Tissue-Tec (Sakura Finetek), frozen in isopentane, and kept at -80°C. Immunofluorescence staining was performed using ATLO GC-derived, RLN GC-derived antibodies, anti-mouse CD11c (N418, Serotec), alpha smooth muscle actin (SMA, 1A4, SIGMA), CXCR4(UMB2, Abcam), RelB (C-19, Santa Cruz Biotechnology), CD35 (8C12, Biosciences), and BCL6 (K112-91, Biosciences). Fresh frozen sections (10 µm) from mice tissues and human carotid plaques were prepared by a cryostat and stored at 80 °C. Tissue sections were heated on the hotplate at 37 °C for 1 min and air dried at RT for 30 min. Sections were fixed with 4% paraformaldehyde (PFA) for 3 min at 4 °C, followed by rinsing with PBS. After diving in acetone for 2 min at RT, the sections were rinsed in PBS for 10 min. Primary antibodies (diluted in PBS with 0.25% BSA) were incubated overnight at 4 °C in a humid chamber. Sections were incubated with fluorescently labeled secondary antibodies (diluted in PBS with 0.25% BSA) and 4’, 6 diamidino 2 phenylindole DAPI (0.1 g/ml) in a humid chamber for 60 min in the dark after washing three times with PBS. After washing with PBS, slides were mounted with Fluoromount G (S3023, DAKO). Human sections were heated on the hot plate at 37 °C for 1 min and air dried at RT for 30 min. Sections were fixed with 4% PFA for 3 min at 4 °C, followed by rinsing with PBS. After fixation, specimens were blocked with blocking buffer (1 X PBS / 5% normal serum / 0.3% Triton X-100) for 30 min at RT, and incubated with diluted ATLOs derived antibodies overnight at 4 °C, and then rinsed with PBS for 3 X 5 min. Specimens were incubated with an antibody without a fluorescent label fluorescent label (purified goat anti mouse IgG, Santa Cruz) for 1 hr at RT. After that, specimens were incubated with secondary antibodies (donkey anti goat, Dianova) with fluorescently and DAPI (1: 500, 0.1 g/ml) for 1 hr at RT, then sections were washed with PBS in the dark and mounted with Fluoromount G. For negative controls, staining was performed without primary antibodies or isotype controls. Stained sections were analyzed using a confocal laser scanning microscope Leica SP8 3X (Mannheim, Germany). All images were prepared as TIF files by imageJ or Leica LAS-X (V1.2) software and exported into Adobe IIIustrator CS6 for figure arrangements.

###### Total cholesterol assay

Blood was collected into a EDTA precoated tube by using 1 ml syringes with a 23-gauge needle anticoagulated blood samples from mice were centrifuged at 6000 rpm for 10 min, then the plasma was collected and used for the determination of total cholesterol according to the manufacturer’s instructions using the cholesterol quantitation kit (Sigma, MAK043-1KT).

###### Statistical analyses

Data were analyzed using GraphPad Prism (version 10) or in R (version 4.2.1). The bar plots were presented as mean ± standard error of mean (SEM). For comparisons of discrete variables, data distribution was tested by Shapiro–Wilk test. For data that followed a Gaussian distribution, student T-test was used to compare difference between two groups or a one-way analysis of variance (ANOVA) with Bonferroni post hoc test was used to perform statistical analysis among multiple groups. For repeated measurement data, such as body weight and antibody levels across multiple periods and among multiple groups, repeated measures two-way ANOVA followed by a post hoc test was performed for analysis. As some datasets did not follow Gaussian distribution, the difference between two groups was compared by two-sided Wilcoxon rank-sum test and difference between three or more groups was analyzed by non-parametric Kruskal–

Wallis H test with Dunn’s post hoc test for pairwise comparisons. For categorical variables, comparisons were formally analyzed by Pearson’s Chi-square test and followed by post hoc pairwise comparisons, which conduct Fisher’s exact test for any of expected frequencies < 5 or Chi-square test for all expected frequencies > 5. The Benjamini–Hochberg procedure was applied to adjust *P* values and control the false discovery rate. Differences were considered significant at a two-tailed *P* value < 0.05. *: *P* <0.05; **: *P* <0.01; ***: *P* <0.001; n.s.: not significant.

###### Data availability

The BCR sequences and associated metadata for each sample, including mouse ID, tissue, genotype, mutation summary, isotype summary, and clonality for Rep-seq, as well as the count matrix and metadata (tissue, cluster, cell type, and BCR-related information) for scRNA-seq data, and IP proteomics data have been deposited in Figshare (10.6084/m9.figshare.27022876). The public GC B cells data used in our study can be downloaded from the GEO database under accession numbers GSE154634.

