## Supplemental Data 1 for "Self-Reactive B Cells in Artery Tertiary Lymphoid Organs Encode Pathogenic High Affinity Autoantibody in Atherosclerosis"

[26]These authors contributed equally
[27]Senior author
[28]Lead contact

**Figure S1- Figure S6**

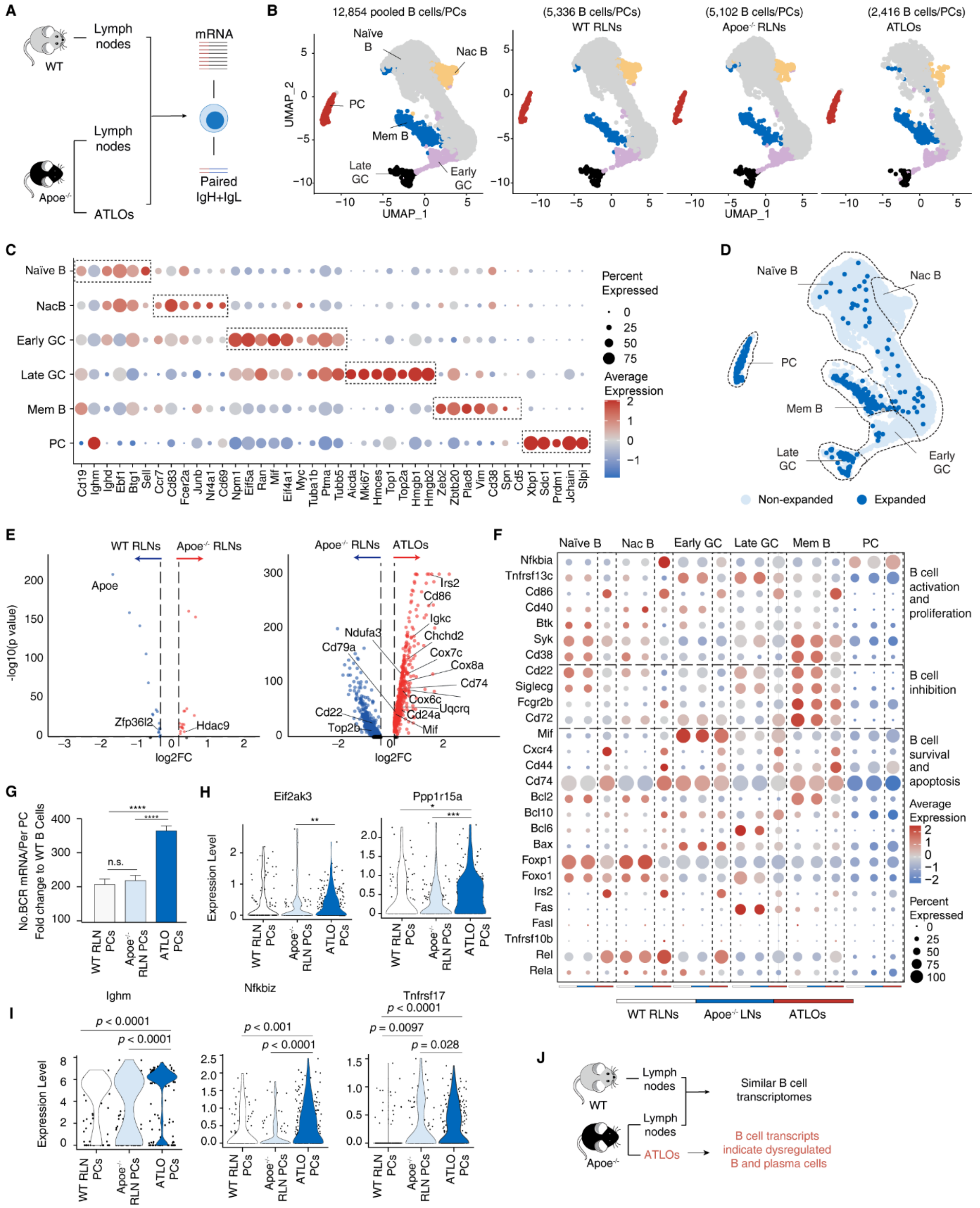


**Figure S1: ScRNA-seq/scBCR-seq pairing reveals tolerance dysfunction in ATLOs**

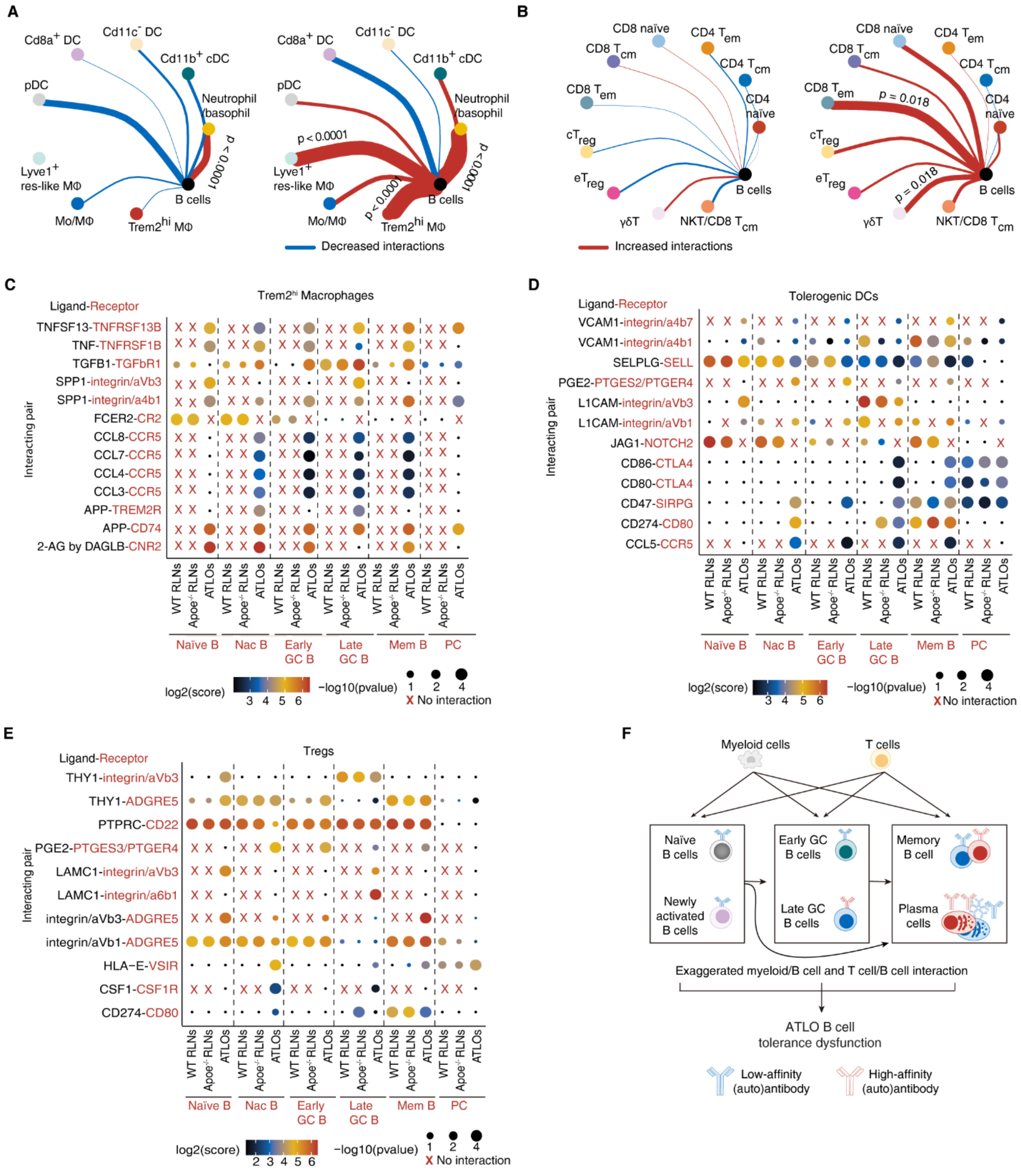


**Figure S2: Modeling of cell-cell interactions reveals aberrant regulation of ATLO B cell immunity**

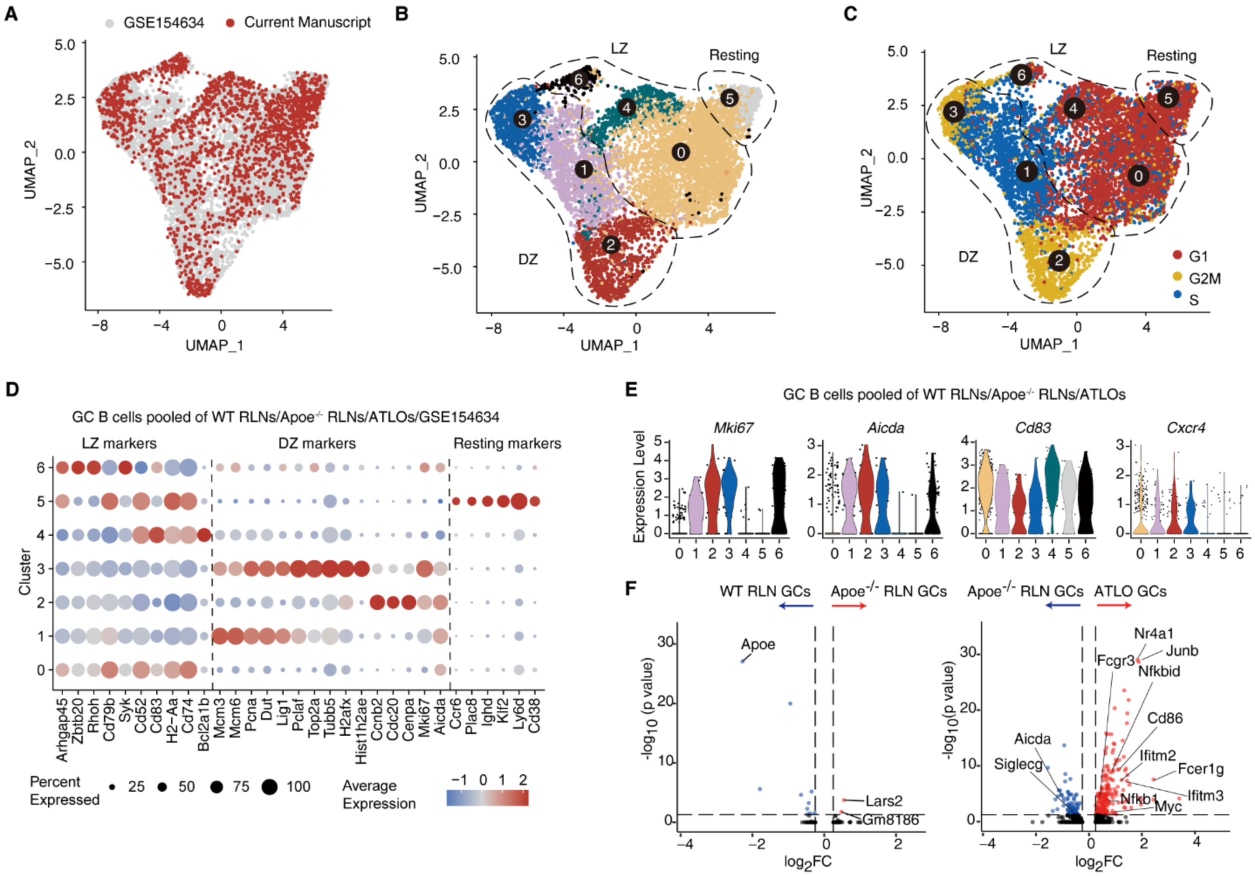


**Figure S3: B cell transcript signatures are dysregulated in ATLO GCs**

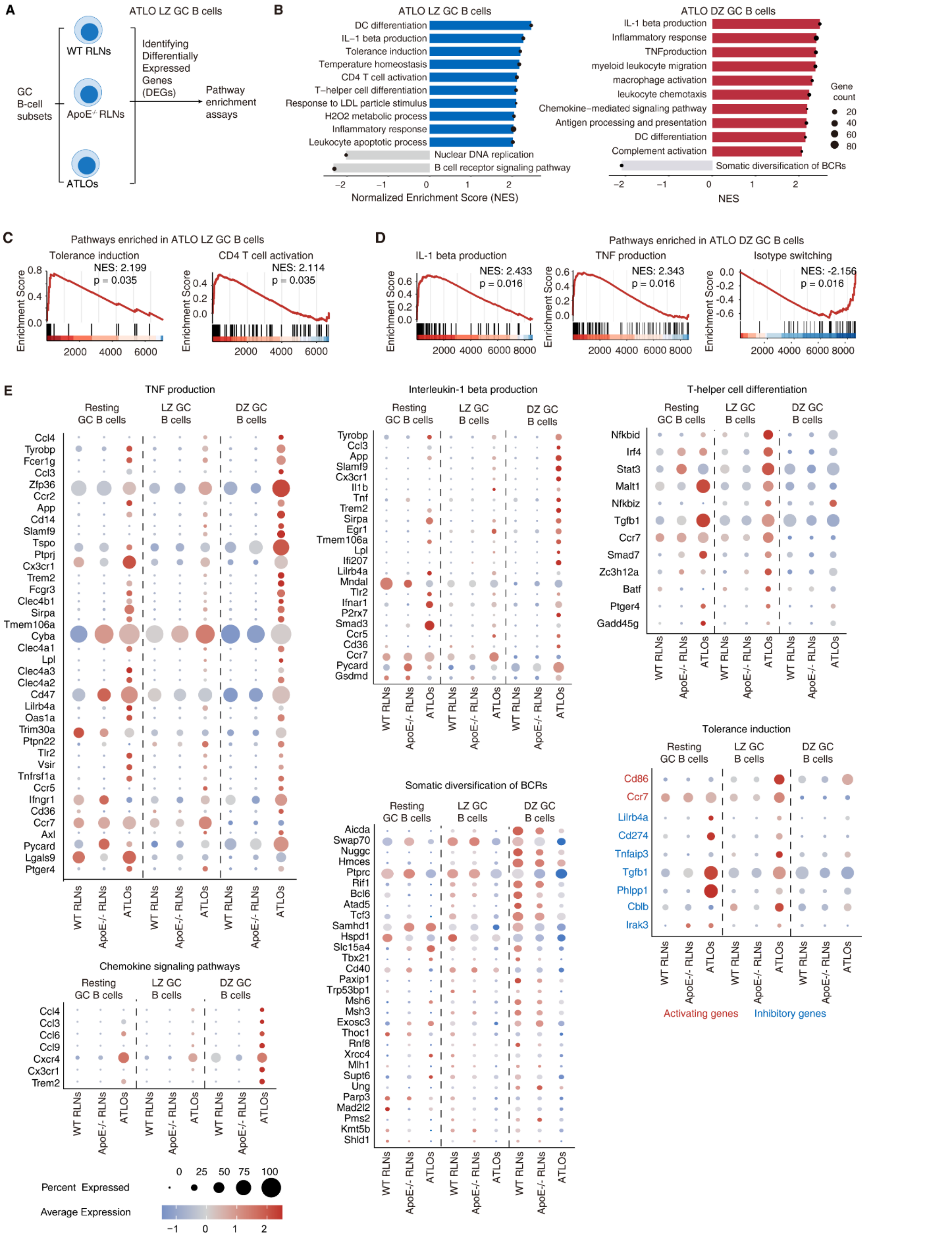


**Figure S4: ATLO GC B cells show increases in GC B cell-associated cytokine pathways and defects of tolerance maintenance**

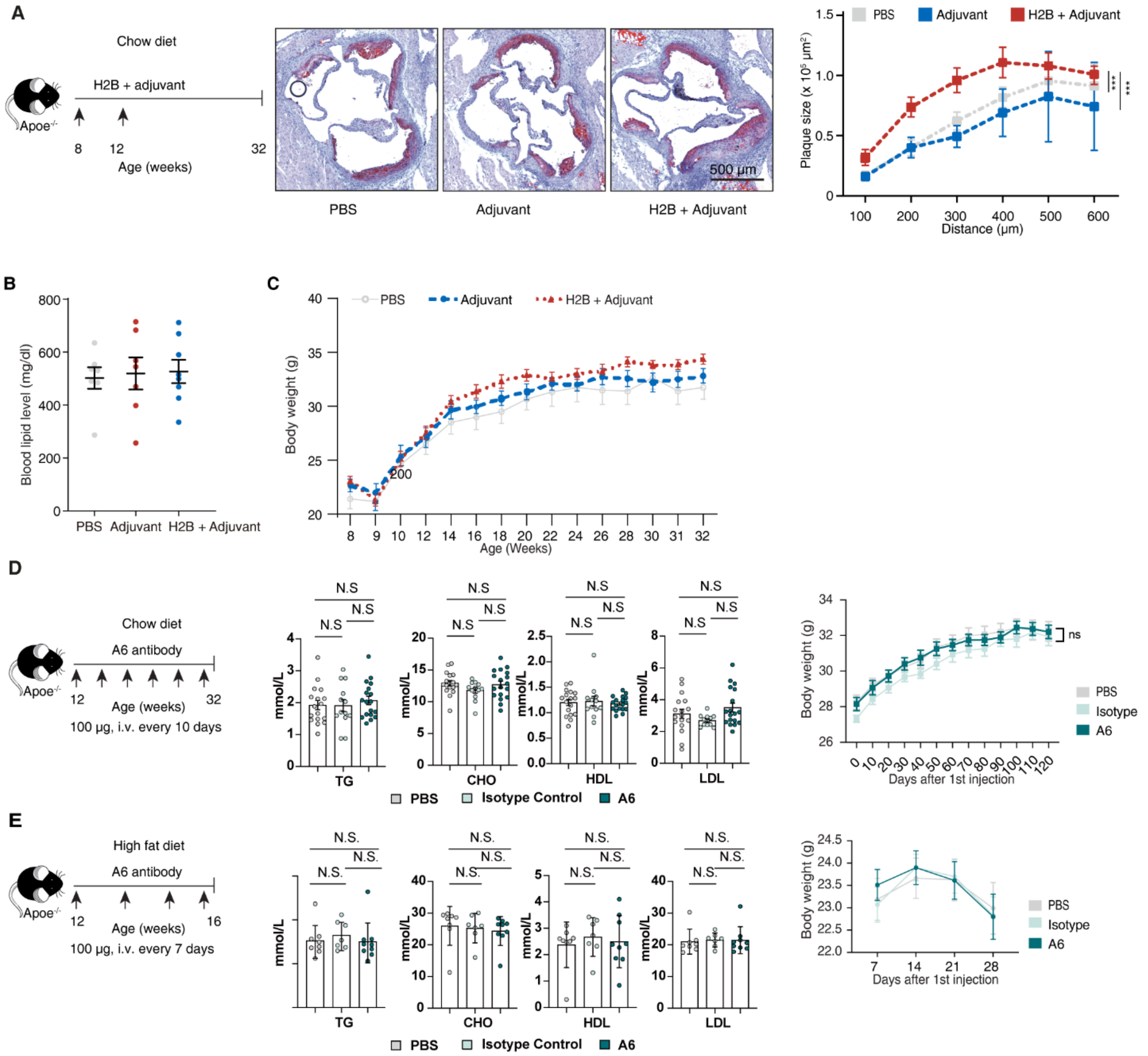


**Figure S5: H2B autoimmune pathway did not impact blood lipids.**

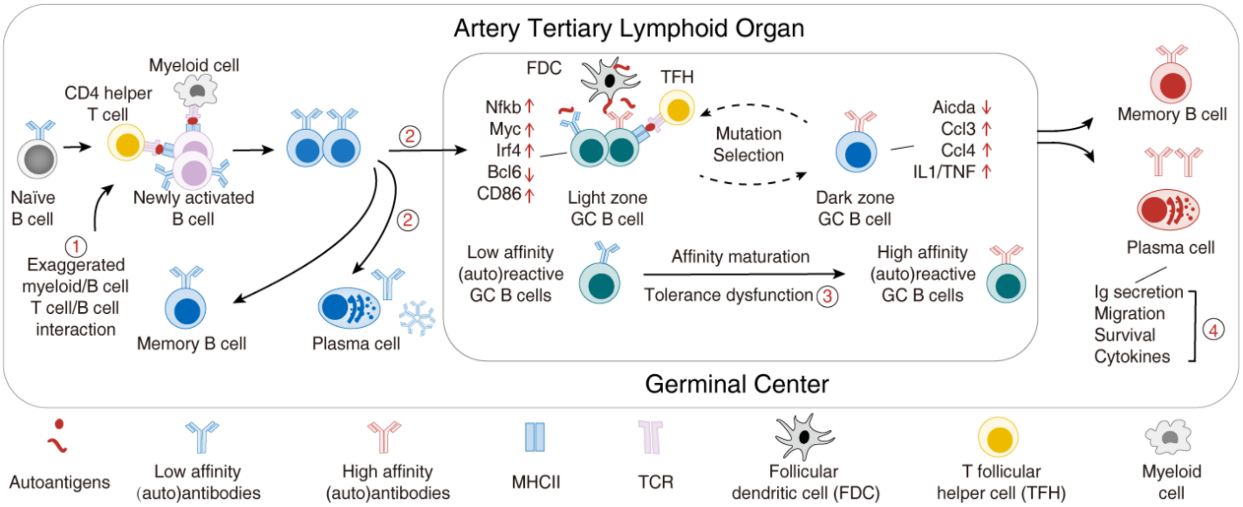


**Figure S6 Hypothetical choreography of the B cell tolerance breakdown in ATLOs**

**Table S1 – Table S2**

**Table S1: Immunoglobulin gene repertoire and reactivity of antibodies from WT and Apoe$^{-/-}$ mice (table S1 provided as separated excel file)**

**Table S2.** Demographic and clinical characteristics of the study cohort (N = 495)

| Category | Characteristic | Value |
|---|---|---|
| **Demographics** | Age, years | 49.8 ± 9.6 |
| | Sex, male | 303 (61.2%) |
| **Anthropometrics** | Body Mass Index (BMI), $kg/m^2$ | 24.0 ± 2.9 |
| | Height, cm | 165.6 ± 7.8 |
| | Weight, kg | 66.4 ± 11.4 |
| **Clinical Measures** | Systolic Blood Pressure, mmHg | 123.7 ± 7.8 |
| | Diastolic Blood Pressure, mmHg | 75.7 ± 10.4 |
| **Laboratory Values** | Total Cholesterol, mmol/L | 5.58 ± 1.11 |
| | LDL Cholesterol, mmol/L | 3.54 ± 0.83 |
| | HDL Cholesterol, mmol/L | 1.47 ± 0.31 |
| | Triglycerides, mmol/L | 1.56 ± 1.06 |
| | eGFR, mL/min/1.73 $m^2$ | 88.1 ± 16.6 |
| | Plasma glucose, mmol/L | 5.15 ± 1.19 |
| | HbA1c, % | 5.57 ± 0.78 |
| | Homocysteine, µmol/L | 11.89 ± 2.89 |
| | Anti-H2B antibody, ng/mL | 788.4 ± 992.5 |

Values are presented as means ± standard deviation (SD) or number (percentage). eGFR, estimated glomerular filtration rate; HbA1c, glycated hemoglobin A1c.
